# The Translatome of Senescent Cells Revealed by Sequencing Actively Translated mRNA

**DOI:** 10.64898/2026.02.27.708233

**Authors:** Maxfield M.G. Kelsey, Radha L. Kalekar, John M. Sedivy

**Affiliations:** Center on the Biology of Aging, Brown University, Providence, Rhode Island 02903, USA; Department of Molecular Biology, Cell Biology and Biochemistry, Brown University, Providence, Rhode Island 02903, USA

**Keywords:** Cellular Senescence, Long Interspersed Nucleotide Elements, Gene Expression Regulation, Codon Usage, Protein Biosynthesis, Aging

## Abstract

Cellular senescence drives aging-related tissue dysfunction through the senescence-associated secretory phenotype (SASP), an inflammatory secretome linked to retrotransposable element (RTE) derepression. Transcriptomic and proteomic approaches have extensively characterized the senescence program, but key gaps remain: transcript abundance is a poor proxy for protein output, limited proteomic depth misses low-abundance proteins, and the highly repetitive sequences of RTEs compromise locus-level peptide attribution. To bridge these gaps, we used AHARIBO (AzidoHomoAlanine-mediated RIBOsome isolation), which captures actively translated full-length mRNAs, to profile the translatome of proliferating, senescent, and late-senescent human fibroblasts. Comparing these ribosome-associated transcripts with the total mRNA pool revealed marked post-transcriptional regulation of key senescence programs. Inflammatory SASP components were translationally depleted in senescence, and these transcripts were enriched for AU-rich element–binding protein motifs, including the ZFP36 family, implicating these proteins in post-transcriptional gating of inflammatory signaling. Transcriptome-wide, translational efficiency was associated with 3’UTR GC content and specific RNA-binding protein and microRNA (miRNA) motifs. We also observed a striking wobble-position codon bias: a proliferation-specific program favoring A/U-ending codons collapsed in senescence, disproportionately affecting cell-cycle and proliferation gene sets. By pairing AHARIBO with a sample-specific reference genome incorporating non-reference L1 insertions, we resolved translation of individual L1 loci and identified two intact L1HS elements with sustained activation in senescence. One of these, L1HS_14q23.2_3, independently identified in multiple experiments, emerges as a candidate intact L1 locus for producing inflammatory cDNA species. These findings implicate translational control as an important regulatory layer shaping the senescent program.

## Introduction

Aging involves the gradual buildup of cellular damage that reduces tissue function and resilience. Among the processes that contribute to age-related decline, cellular senescence has emerged as an important driver of chronic inflammation, tissue dysfunction, and degenerative diseases (Campisi, 2013; Lopez-Otin, Blasco, Partridge, Serrano, & Kroemer, 2013). Senescent cells accumulate in aged tissues and increase in response to telomere attrition, genotoxic stress, and oncogenic signaling (van Deursen, 2014). Although senescent cells no longer divide, they remain metabolically active and influence the surrounding tissue.

A central effector of senescent cell-mediated tissue dysfunction is the senescence-associated secretory phenotype (SASP) (Campisi, 2013), a diverse set of cytokines, chemokines, growth factors, and proteases that promote inflammation and disrupt tissue homeostasis during aging. Genetic and pharmacological studies demonstrate that reducing senescent cell burden or attenuating the SASP improves tissue function and extends healthspan in animal models (Baker et al., 2011; Xu et al., 2018). However, elimination of senescent cells can impair tissue repair and regeneration in certain contexts (Demaria et al., 2014), highlighting the need to better understand how different aspects of the senescence program are regulated.

Senescent cells exhibit a global decline in protein synthesis, driven by reduced ribosome biogenesis, altered mTOR signaling, and activation of cellular stress pathways (Hernandez-Segura et al., 2017; Laplante & Sabatini, 2012). Yet against this backdrop of broad translational repression, certain transcripts continue to be efficiently translated (Li et al., 2013), implying that senescent cells operate under heightened translational selectivity. This selectivity is likely shaped by a combination of regulatory inputs, including RNA-binding proteins, miRNAs, and intrinsic RNA sequence features and secondary structures (Biziaev et al., 2024; Courel et al., 2019; Dong, Wei, Zhang, & Wang, 2018; Srikantan, Marasa, Becker, Gorospe, & Abdelmohsen, 2011). A direct consequence of such selectivity is that mRNA abundance becomes a poor proxy for protein output, as borne out by the weak correlation between transcriptomic and proteomic measurements in senescent cells (Sullivan et al., 2021; Takemon et al., 2021). Resolving how the senescent program is regulated therefore requires methods that directly assess translation.

Ribosome profiling (Ribo-seq) is a widely used method for defining translational landscapes, but its application to replicative senescence faces some limitations. Ribo-seq captures ribosome-protected fragments irrespective of whether the ribosomes are actively elongating, transiently paused, or stably stalled. In senescent cells, where translational stress and ribosome stalling are prevalent (Zhou et al., 2025), ribosome occupancy does not necessarily reflect protein synthesis — indeed, studies of stress and unfolded protein response signaling have shown that ribosomes remain associated with mRNAs even as translation is globally repressed (B. Liu, Han, & Qian, 2013; T. Y. Liu et al., 2017). The large input requirements of Ribo-seq further limit its utility in senescence models with reduced proliferative capacity.

These limitations are particularly acute for studying the expression of RTEs such as the long interspersed nuclear element 1 (LINE-1, or L1) (De Cecco et al., 2019; Simon et al., 2019). L1 expression increases in senescence and contributes to genomic instability and to chronic inflammation via cytoplasmic nucleic acid sensing of cDNAs generated by the L1-encoded ORF2 reverse transcriptase (De Cecco et al., 2019). Although most L1 transcripts arise from truncated or non-coding copies, the detection of L1 proteins and cDNAs in senescent cells necessitates that some loci be productively translated (Ingolia, 2014; Munot et al., 2025). However, identifying which loci produce L1 proteins (ORF1 and ORF2) has remained elusive. RNA sequences generated by Ribo-seq are too short for unique assignment to the highly repetitive L1 loci, and proteomic methods cannot resolve L1 expression at the locus level because ORF1 and ORF2 peptides are largely identical across evolutionarily recent intact L1 loci.

## Results

### The translatome uncouples from the transcriptome in senescence

To profile actively translated mRNAs across proliferating (PRO), senescent (SEN), and late senescent (LSEN) primary human fibroblasts (LF1), we applied AHARIBO (Fig. 1A), a method that selectively enriches for elongating ribosomes by metabolically labeling nascent peptides with the methionine analog azidohomoalanine, whose azide side chain enables capture of the associated ribosome–mRNA complexes via Click-iT chemistry (Minati et al., 2021). Comparing AHARIBO profiles with total mRNA-seq from matched samples allowed us to disentangle transcriptional from translational regulation, and pairing this approach with a sample-specific reference genome (Dalgarno et al., 2025) enabled locus-level resolution of L1 translation in senescent cells. We refer to RNA-seq of total mRNA as the “transcriptome” and AHARIBO RNA-seq as the “translatome”.

**Fig. 1.**
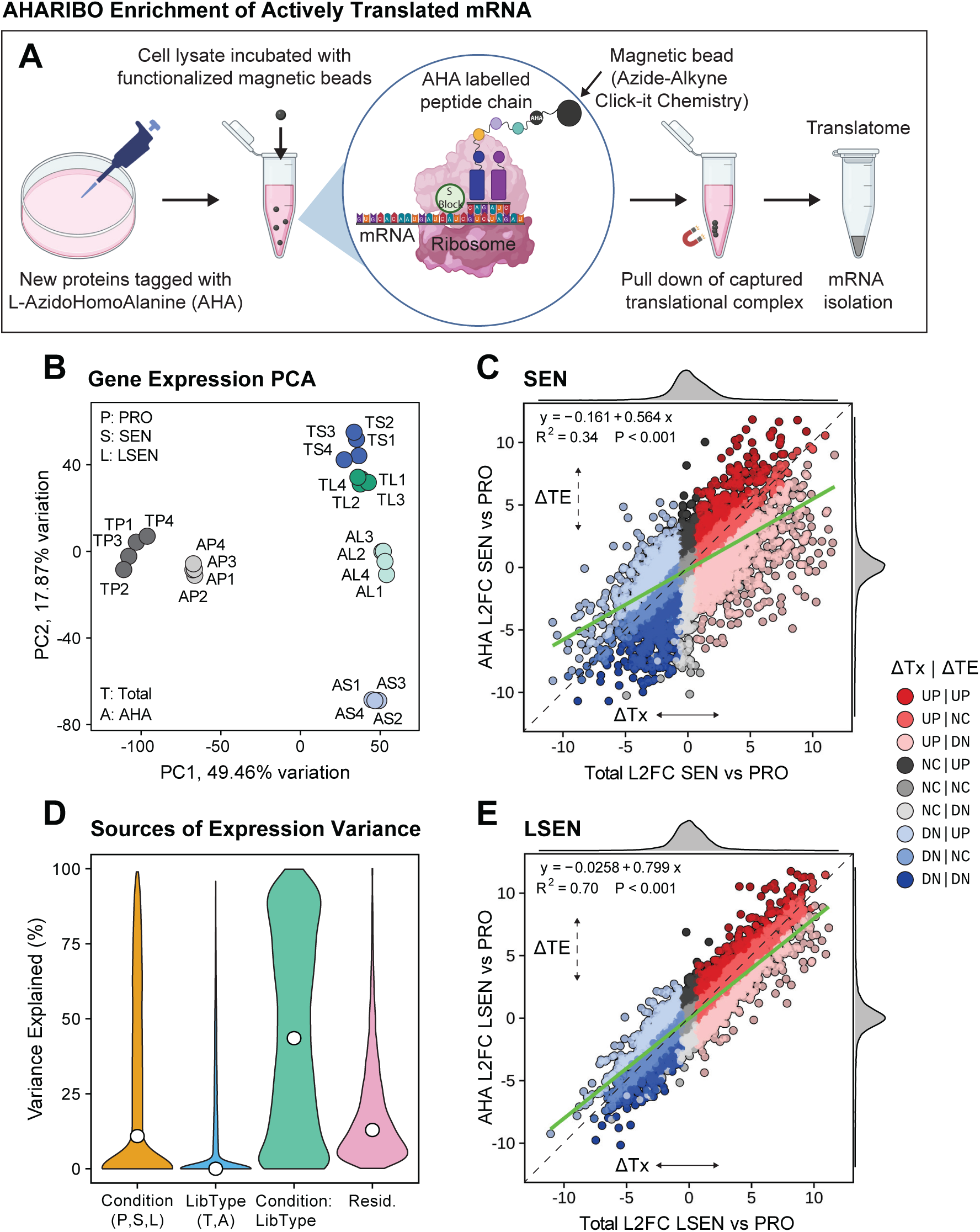
AHARIBO profiling enriches for the active translatome. **A** Diagram of the AHARIBO method used to enrich for actively translated transcripts. **B** Principal Component Analysis (PCA) of gene expression profiles. **C,E** Scatter plots showing expression changes in the translatome vs. the transcriptome for (C) SEN vs. PRO and (E) LSEN vs. PRO. Deviations from the diagonal denote changes in translational efficiency, ΔTE. We fit a linear model (green line) and report its parameters and statistics. We classified genes into distinct regulatory modules (see Fig. S1B & Methods) based on the directionality of changes in their transcription (ΔTx) and in their translational efficiency (ΔTE). Genes are colored by their regulatory module. DN, down; NC, no change. **D** Violin plots of the variance in gene expression explained by various factors, for all expressed coding genes (n = 13,583; average ≥100 counts across conditions). White circle shows the median. Expression was modeled with linear mixed-effects models (random intercepts for biological condition, technical condition, and their interaction) using the variancePartition package[46]. Yellow: biological condition; blue: library preparation method; green: interaction term for biological condition and library preparation method; pink: model residuals (unexplained variance).

Principal component analysis revealed that samples clustered by senescence state along PC1 and by library type along PC2, with the SEN translatome most distinct from its matched transcriptome (Fig. 1B). To characterize the translational regulation of gene expression in senescence, we plotted translatomic against transcriptomic gene expression changes in SEN and LSEN (Fig. 1C,E). Genes falling on the diagonal change equivalently in both pools, while off-diagonal genes are translated more (above the diagonal) or less (below the diagonal) than their mRNA levels would predict; we refer to this divergence as a change in translational efficiency (ΔTE). SEN cells exhibited a more pronounced uncoupling of the translatome from the transcriptome than LSEN (SEN R² = 0.34, Fig. 1C; LSEN R² = 0.70, Fig. 1E). Transcriptomic changes in SEN were largely opposed by changes in translational efficiency, as evidenced by a flattened slope (green line, Fig. 1C). This indicates that SEN is substantially post-transcriptionally regulated and characterized by extensive translational buffering, while these effects are attenuated in LSEN.

Quantifying these effects across all conditions simultaneously, we fit a multi-level linear model and partitioned each gene’s expression variance among biological condition (PRO vs. SEN vs. LSEN), library type (total mRNA vs. AHARIBO), and their interaction (Fig. 1D). Variance attributable to biological condition alone reflects regulation shared by the transcriptome and translatome, whereas variance attributable to the interaction term reflects changes in the translatome that are not simply inherited from the underlying mRNA level. For most genes, the interaction term explained the largest share of variance, confirming that senescence reshapes the translatome beyond what is predicted by transcriptional changes alone. These changes in translational efficiency must be interpreted in the context of a global decrease in translation in senescence. This decrease has been reported previously (Payea et al., 2024) and is evidenced in our data by the systematic downregulation of ribosomal proteins (Fig. S1A). Hence, genes with a +ΔTE may nevertheless be translated at a lower absolute level in senescence owing to the global decrease in translation.

### Translational efficiency regulates canonical and non-canonical senescence programs

We classified genes into distinct regulatory modules (Fig. 2A, S1B) based on the direction of changes in their transcription (ΔTx) and translational efficiency (ΔTE), and visualized these modules by hierarchical clustering. We considered genes with translatomic changes that differed by >2-fold from their transcriptomic changes to be post-transcriptionally regulated in senescence. A somewhat larger proportion of genes fell into translationally affected classes in SEN (12,349) than in LSEN (11,049), and a greater fraction of SEN translationally affected genes exhibited discordant changes (e.g., ΔTx UP, ΔTE DOWN, 6,112/12,349) than in LSEN (5,047/11,049).

**Fig. 2.**
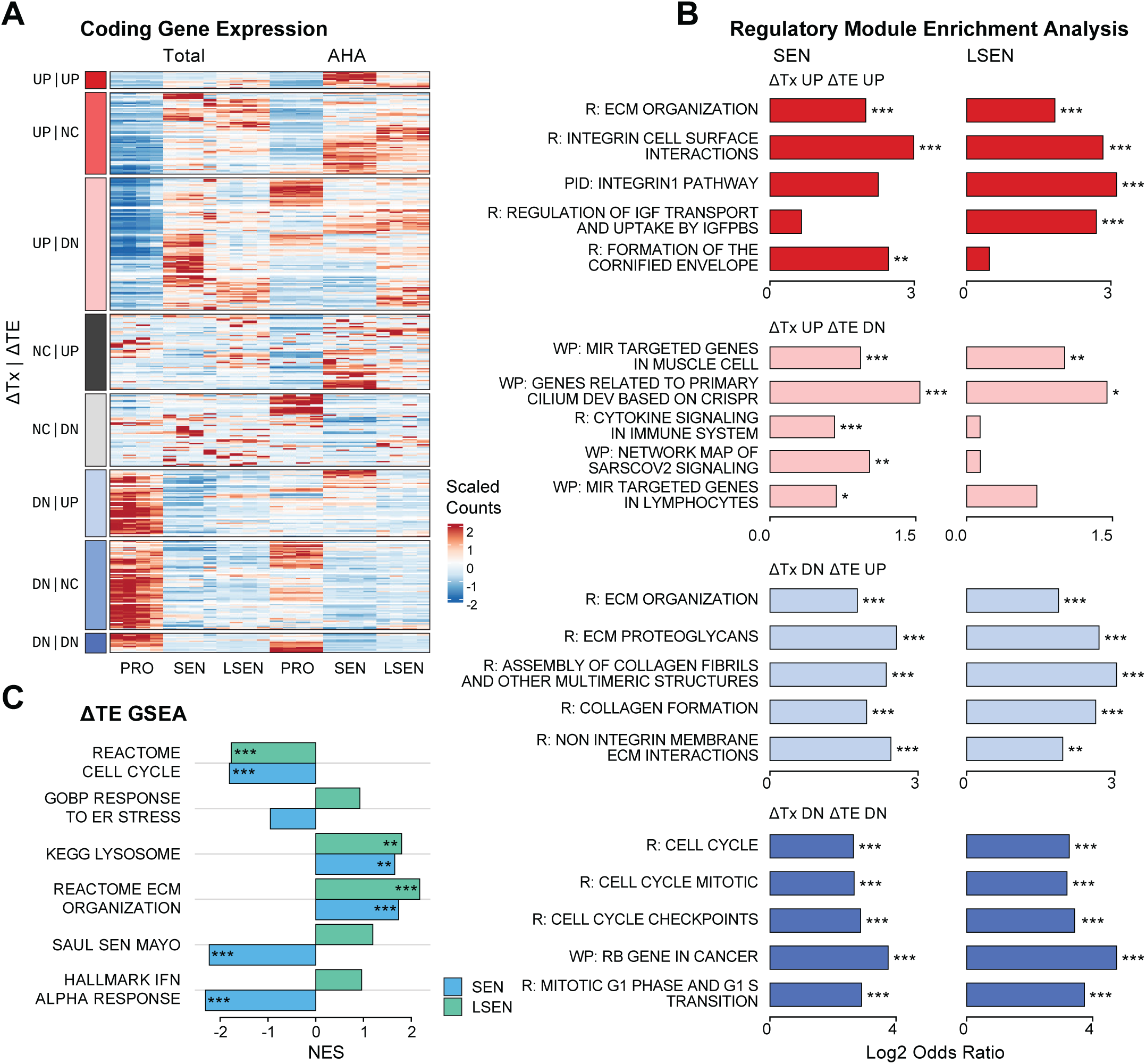
The translatomic landscape of senescence reveals extensive and stage-dependent translational regulation. **A** Heatmap of gene expression across all samples, grouped by regulatory module. **B** Over-representation analysis (ORA) of selected regulatory modules in MSigDB Canonical Pathways (CP). Statistical significance was assessed using Fisher’s exact test, and FDR-adjusted p-values are reported. **C** Gene Set Enrichment Analysis (GSEA) of ΔTE ranking, showing Normalized Enrichment Scores (NES) for key senescence-associated pathways in SEN (blue) and LSEN (green).

We next performed a targeted analysis of selected senescence-associated gene sets to determine whether they exhibited signs of translational regulation. We performed GSEA on TE-ranked genes (Fig. 2C) and a regulatory module overrepresentation analysis (Fig. S1C). The Reactome Cell Cycle gene set was transcriptionally and translationally depleted in both SEN and LSEN, and strongly enriched in the “ΔTx DOWN ΔTE DOWN” module. In contrast, KEGG Lysosome and Reactome extracellular matrix (ECM) organization gene sets were both transcriptionally and translationally enriched in both senescence conditions. Surprisingly, the SenMayo gene set (Saul et al., 2022) (largely composed of inflammatory SASP factors) and the Hallmark Interferon-Alpha Response gene set were transcriptionally enriched but translationally depleted in SEN, and this depletion was relieved in LSEN.

To interpret this depletion, we considered that genes can have decreased translational efficiency regardless of whether their absolute representation in the translatome increases or decreases: they can exhibit (i) a transcriptional increase but no translational increase, (ii) no transcriptional increase but a translational decrease, or some combination of (i) and (ii). We therefore examined whether, despite having a negative ΔTE, SenMayo and IFN-alpha response genes were upregulated in the SEN vs. PRO translatome. SenMayo was slightly upregulated, while IFN-alpha response genes were downregulated in SEN (Fig. S1D,E). Both were sharply higher in LSEN (Fig. S1D,E). This suggests that inflammatory genes in SEN are initially transcriptionally upregulated, but that this upregulation is transiently opposed by translational downregulation, which is subsequently relieved.

We next performed an unbiased analysis of the MSigDB Canonical Pathways gene sets (Liberzon et al.) to determine which were most enriched in our regulatory modules (Fig. 2B). This analysis corroborated our targeted analysis of senescence-associated pathways. The top “ΔTE UP” pathways pointed to sustained ECM and cytoskeletal changes in senescence. The top “ΔTx DOWN ΔTE DOWN” pathways were cell-cycle-related, and several top “ΔTx UP ΔTE DOWN” pathways were immune-related. Beyond the canonical senescence-associated pathways noted above, this analysis pointed us to a pathway we had not previously investigated in the context of senescence. One of the top “ΔTx UP ΔTE UP” pathways in SEN was Formation of the Cornified Envelope (CE), a tough cross-linked protein structure that typically forms under the cell membrane in terminally differentiated keratinocytes during cornification, a unique form of programmed cell death. We prioritized this hit because the structure is not known to form in fibroblasts (Candi, Schmidt, & Melino, 2005), yet several core CE components—loricrin, involucrin, filaggrin, desmoplakin, annexin A1, and keratin 14—were actively translated in senescent cells despite being absent in proliferating ones, alongside upregulation of transglutaminase 2, which can functionally substitute for the epidermal cross-linking enzyme transglutaminase 1 (Fig. S2B,D). This may represent either co-option of cornification machinery to reinforce the cytoskeleton and contribute to the stiffened, flattened senescent morphology, or non-programmed cell identity drift.

Taken together, these data establish that the senescence program is not merely a transcriptional state, but that key senescence programs are heavily influenced by stage-specific translational control mechanisms.

### The SASP is translationally repressed in senescence and derepressed in late senescence

We next sought a detailed picture of the translational regulation of SASP genes by examining the secreted subset of SenMayo genes (Saul et al., 2022). We first plotted translatomic vs. transcriptomic changes for SASP genes (Fig. 3A, S3A), which revealed that SASP components were slightly upregulated in the SEN translatome compared to PRO, despite a profound decrease in their translational efficiency (Fig. 3A). Because SASP proteins are secreted and translated through the ER membrane, we sought to confirm that capture efficiency for such transcripts was not compromised in AHA, comparing the ΔTE of SASP genes and of genes encoding other secreted factors (GOBP Protein Localization to Extracellular Region gene set) to that of all coding genes. This analysis showed that the SASP is translationally depleted in SEN (p < 0.001 for both comparisons) but not in LSEN (Fig. 3B), and that ER-translated transcripts are not systematically depleted in AHA, lending confidence that the observed depletion of SASP transcripts is not a technical artifact.

**Fig. 3.**
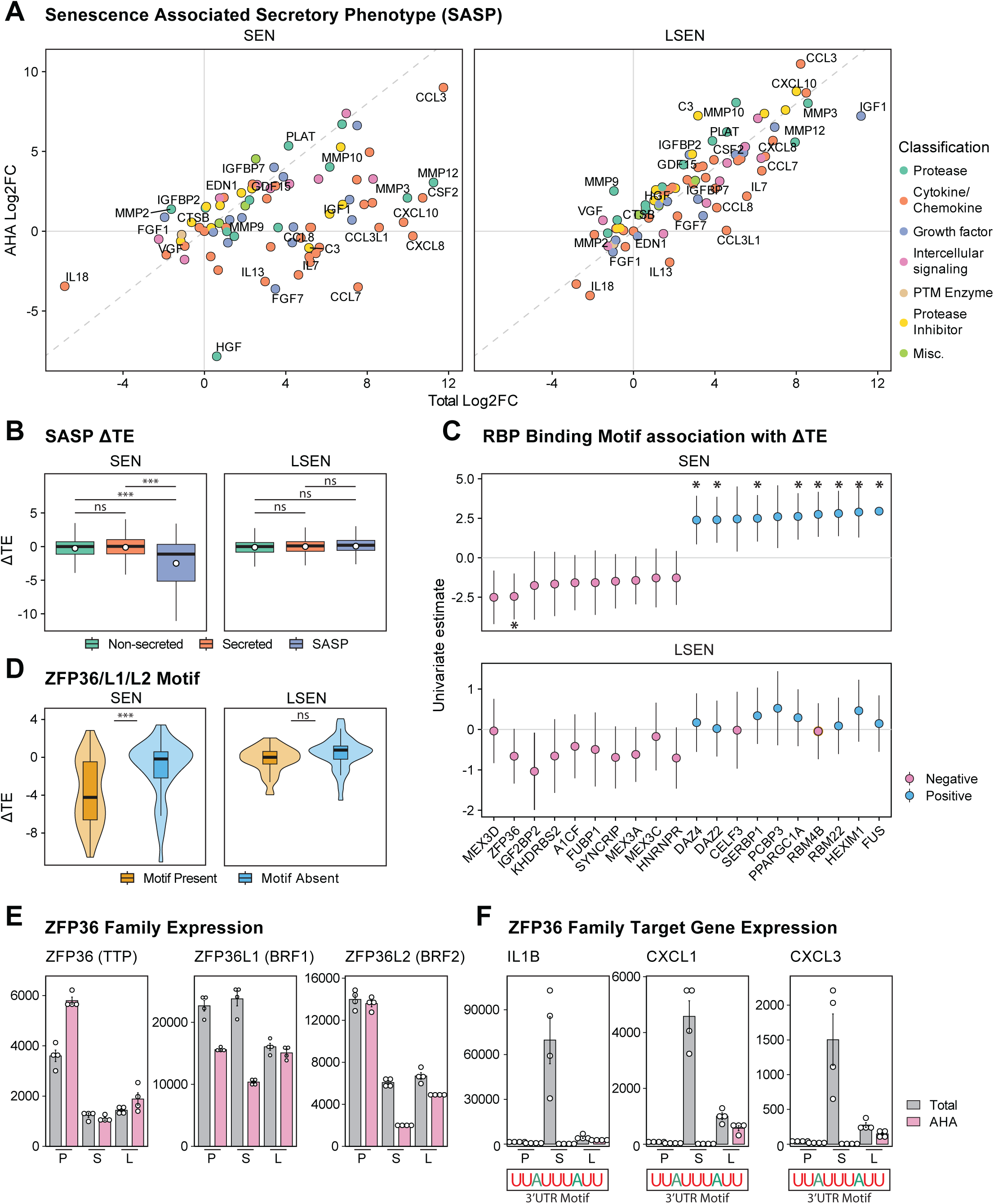
The senescence-associated secretory phenotype (SASP) is translationally repressed in senescence and derepressed in late senescence. **A** Scatter plots comparing the expression changes of SenMayo SASP genes in the translatome vs. the transcriptome for SEN vs. PRO (left) and LSEN vs. PRO (right). Genes are colored by their SenMayo classification. **B** Boxplot of ΔTE for SASP, non-SASP secreted, and non-SASP non-secreted coding genes. Statistical significance was assessed using a two-sided Wilcoxon rank-sum test. **C** UTR sequences of SenMayo SASP genes were scanned for RNA-binding protein motifs from the CIS-BP database using Gimme Motifs. For each RBP, ΔTE was regressed on binary motif presence using a univariate linear model. Point estimates and 95% confidence intervals for the top RBP’s effect on ΔTE are shown. FDR-adjusted p-values are denoted by: p ≤ 0.05 *. **D** Violin plot of ΔTE for all SASP genes stratified by ZFP36/L1/2 binding motif presence. Statistical significance was assessed using a two-sided Wilcoxon rank-sum test. **E,F** Gene expression plots for ZFP36/L1/2 (E) and selected SASP genes (F) with ZFP36/L1/2 binding motifs. Error bars represent mean ± SEM. Statistical significance was assessed using DESeq2 FDR-adjusted p-values; p ≤ 0.05 *, ≤ 0.01 **, and ≤ 0.001 ***.

To identify candidate post-transcriptional regulators of the SASP, we performed univariate linear regression and elastic net regression of ΔTE against RBP motif presence in SASP transcript UTRs (Fig. 3C, S3E,F). The top factors associated with reduced ΔTE were enriched for AU-rich element (ARE)-binding proteins, including the ZFP36 family (ZFP36/TTP, ZFP36L1/BRF1, ZFP36L2/BRF2), MEX3A/C/D, A1CF, and SYNCRIP, with effect sizes substantially larger in SEN than in LSEN, consistent with post-transcriptional regulation of the SASP being concentrated in the former. That the ZFP36 family emerged as the top predictor is concordant with the established role of ZFP36L1 in SASP regulation in oncogene-induced senescence (Herranz et al., 2015). ZFP36/ZFP36L1/ZFP36L2 bind AREs in 3′UTRs to drive both translational silencing and CCR4-NOT-mediated decay (Otsuka et al., 2020; Rasch, Weber, Izaurralde, & Igreja, 2020; Tollenaere et al., 2019); here, they were themselves downregulated in senescence (Fig. 3E), yet motif-bearing SASP transcripts accumulated at the mRNA level while being translationally repressed in SEN (Fig. 3A, 3D), pointing to a silencing activity decoupled from decay. This translational repression of motif-bearing SASP transcripts was relieved in LSEN, with partial re-engagement of decay (Fig. S3C,D), supporting a model in which reduced ZFP36/ZFP36L1/ZFP36L2 in SEN preferentially suppresses translation of these SASP transcripts, including IL1B, CXCL1, and CXCL3 (Fig. 3F), without driving their turnover. Nevertheless, the highly overlapping AU-rich consensus motifs across the identified RBPs produce strong inter-factor correlations in transcript binding (Fig. S3G), making it difficult to attribute regulation of any given SASP target to a specific factor.

### Young LINE-1 elements are translated in senescence

We next interrogated the translatome for retrotransposable element (RTE) representation. Because the protein products of most coding RTE loci are nearly identical, locus-level assignment of protein/peptide expression is limited. Instead, RNA-seq of translationally active RTE transcripts allowed us to examine which expressed RTE loci produce protein products. Plotting the translational efficiency of young full-length RTEs revealed that L1s were the most translationally engaged family (Fig. 4A). This is consistent with the coding landscape of human RTEs: Alu and SVA elements are non-coding, ERVs are largely too diverged to translate, and only young L1s retain intact open reading frames.

**Fig. 4.**
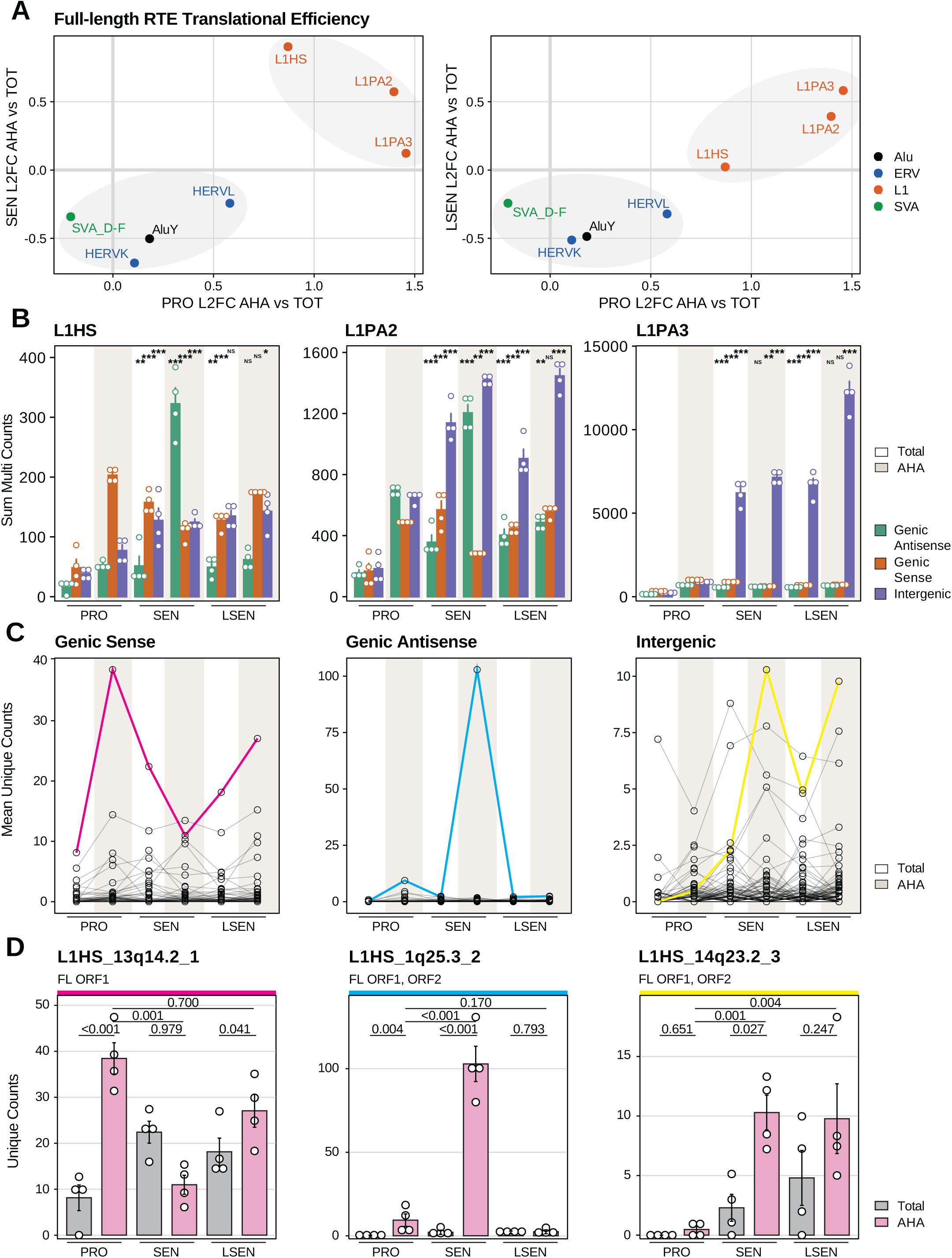
Translational activation of young LINE-1 retrotransposons in senescence. **A** Scatter plots comparing translational efficiency (Log2 fold-change of AHARIBO vs. Total RNA) among full-length, non-genic-sense retrotransposable elements (RTE), grouped by subfamily and colored by family. We show SEN (left) and LSEN (right) vs. proliferating (PRO) baselines. Young LINE-1 subfamilies (L1HS, L1PA2, L1PA3; orange) exhibit high TE relative to other families (Alu, ERV, SVA). Gray ellipses are visual guides grouping young L1 subfamilies and non-L1 RTEs; they are not statistical fits. **B** Aggregate expression (sum of multi-mapping counts) of full-length members of the three youngest L1 subfamilies, stratified by genomic context: genic sense (intronic), genic antisense, and intergenic. Statistical comparisons match each condition to its proliferating counterpart and are based on negative binomial modeling of raw aggregated counts, incorporating DESeq2-derived size factors as a model offset; p-values are FDR-adjusted for all pairwise comparisons in the panel. FDR-adjusted p ≤ 0.05 *, ≤ 0.01 **, and ≤ 0.001 ***. **C** Full-length L1HS locus expression (mean unique counts). Individual lines represent distinct loci; colored lines highlight representative elements for genic sense (magenta), genic antisense (blue), and intergenic (yellow) contexts. **D** Unique-count expression profiles for the three representative FL L1HS loci highlighted in (C), confirming locus-specific translational upregulation. Error bars represent mean ± SEM. DESeq2 FDR-adjusted p-values for comparisons between AHA and TOT are shown for each senescence condition, as well as for the AHA SEN/LSEN vs. PRO comparisons.

We stratified the three youngest L1 subfamilies (L1HS, L1PA2, and L1PA3) by their genomic context: genic sense (intronic), genic antisense, and intergenic (Fig. 4B). Although genic sense elements displayed high counts, their expression often reflects misattributed intronic signal from host-gene transcription, confounding identification of autonomous L1 activity. In contrast, genic antisense and intergenic elements are less likely to register expression as a byproduct of host-gene read-through. Both classes showed marked translational upregulation in senescence relative to proliferating controls (Fig. 4B). This upregulation was dynamic: genic antisense signal was particularly high in the SEN translatome, while intergenic elements showed sustained accumulation into late senescence.

To identify translationally active L1HS loci, we plotted the expression of all full-length (FL) L1HS loci (Fig. 4C). This revealed substantial heterogeneity. Although many loci remained silent, a subset of FL elements exhibited sharp, stage-specific translation. We highlighted one robustly translated element from each genomic context to illustrate the range of observed behaviors (Fig. 4D). The genic sense element L1HS_13q14.2 (intronic) tracked the general translational profile of its host gene, RB1 (Fig. 4D, S4A), consistent with passive read-through from host transcription rather than independent L1 promoter activity. The genic antisense element L1HS_1q25.3 showed a sharp, transient peak of translation confined to SEN, falling back to near-baseline in LSEN. The intergenic element L1HS_14q23.2_3 was upregulated in both the transcriptome and translatome of senescent cells relative to proliferating cells, sustaining elevated expression across both SEN and LSEN.

L1HS_14q23.2_3 retains intact ORF1 and ORF2 coding capacity, a property shared by only a fraction of full-length L1HS loci. These intact loci are of particular interest because they are the source of the inflammatory and genome-destabilizing activities attributed to L1 in senescence, most notably the reverse transcriptase activity of ORF2, which can promote cGAS–STING activation and an IFN-I response. Across the full set of intact, full-length L1HS loci, only two reached significance for upregulation in the AHA translatome at both SEN and LSEN: L1HS_14q23.2_3 and the non-reference element L1HS_3q12.1_1 (Fig. S4C, Fig. S5).

Beyond its sustained translational upregulation, several features set L1HS_14q23.2_3 apart from the broader pool of intact, full-length L1HS elements. In parallel work from our group profiling the 3D genome of late-passage replicatively senescent LF1 fibroblasts by high-resolution Hi-C, total RNA-seq, and Nanopore long-read sequencing, this locus again emerged as one of just two intact L1HS elements meeting genome-wide significance for upregulation in senescence (Dalgarno et al., 2025). It was distinguished from other intact L1HS elements by its chromatin context: the locus simultaneously shifted toward the active A compartment, underwent one of the strongest activating subcompartment transitions in the dataset (B.1.2 to A.1.2), gained a senescence-specific chromatin loop placing it in contact with two upregulated genes, and exhibited a profound loss of CpG methylation at the start of its internal promoter region. L1HS_14q23.2_3 also scored as significantly upregulated in another independent dataset from our group profiling DNA damage–induced senescence in human astrocytes (Woodham, Kelsey, & Sedivy, 2026), indicating that its derepression is not specific to fibroblast replicative senescence or to a single senescence-inducing stimulus, but recurs across cell types and triggers. The sustained transcriptional and translational upregulation of L1HS_14q23.2_3 observed here—in an independent LF1 senescence time course profiled with a different methodology—favors the conclusion that this locus is not merely transcriptionally derepressed but is actively loaded onto translating ribosomes. This makes it a strong candidate among full-length, retrotransposition-competent L1s for contributing to the cytosolic ORF2-derived RT activity implicated in senescence-associated inflammation.

Although L1HS represents the youngest, most retrotransposition-competent L1 lineage in the human genome, locus-resolved translation also revealed activity among slightly older L1 subfamilies. Within L1PA3, one intergenic element stood out: L1PA3_22p13_1 was not expressed in proliferating cells but showed very high transcriptomic and translatomic expression in both SEN and LSEN (Fig. S4B). It lies in a newly resolved subtelomeric region of the human genome (present in T2T-HS1 but absent from hg38, chr22 position 368,301). L1PA3_22p13_1 has partial ORF1 and ORF2 reading frames, the latter of which comprises the entire endonuclease (EN) domain. This EN domain retains catalytic histidine 230, which is responsible for nicking DNA during target-primed reverse transcription (TPRT). Furthermore, truncated ORF2 variants can have enhanced EN activity (Kines et al., 2016). L1PA3_22p13_1 may therefore be a particularly genotoxic element in senescence.

Together, these data confirm that the senescent translatome supports dynamic, autonomous translation of specific, full-length young L1 elements, and nominate L1HS_14q23.2_3 as a likely source of L1 protein products in senescence.

### Sequence features associated with translational remodeling in senescence

To investigate the molecular mechanisms governing translational remodeling in senescence, we searched for sequence features associated with changes in translational efficiency. We first asked whether a shift between cap-dependent and cap-independent translation could account for these changes. The integrated stress response (ISR) represses cap-dependent translation and favors transcripts bearing internal ribosome entry sites (IRES); although eIF2α phosphorylation is elevated in senescence, the ISR is reportedly not activated (Payea et al., 2024). Consistent with this, ΔTE was unaffected by IRES presence in either SEN or LSEN (Fig. S6C), arguing against a switch to cap-independent translation, in agreement with Payea et al. (Payea et al., 2024).

Having ruled out a global change in cap-dependence, we turned to sequence features that could direct selective translation. Examining UTR base composition, we found a substantial positive correlation between 3’UTR GC content and TE in both SEN (R² = 0.10, Fig. S6A) and LSEN (R² = 0.11, Fig. S6B), whereas 5’UTR GC content had negligible influence (R² = 0.01). This asymmetry points to the 3’UTR as a primary locus of sequence-based regulation, consistent with the dominant role of 3’UTR-binding RBPs and miRNAs in translational control.

We next examined trans-acting factors. miRNAs, which predominantly repress their targets (Jonas & Izaurralde, 2015), were strongly skewed toward translational depletion: GSEA recovered 1,002 miRNA-regulated gene sets negatively associated with ΔTE vs. only 70 positively associated (FDR < 0.05; Fig. S6D), consistent with increased miRNA-mediated regulation in senescence. RBP motif enrichment showed the same directionality, with 139 RBPs enriched in translationally depleted transcripts (ΔTE ≤ −2) vs. 38 in enhanced transcripts (ΔTE ≥ 2; Fig. S6F,G). The top depletion-associated motifs were largely shared between SEN and LSEN and survived GC correction, indicating a robust association with translational repression; the two top enhancement-associated motifs, by contrast, were lost after GC correction, leaving it unclear whether they reflect genuine regulation or the underlying GC bias. The fraction of transcripts carrying a motif scaled inversely with motif length, from the short RBM34 motif (∼35-40% of - TE transcripts) to the longer MEX3-family motifs (∼4-5%). No motifs were significantly enriched among transcripts with differential regulation (ΔΔTE) between SEN and LSEN (Fig. S7B).

Taken together, these analyses indicate that the senescent translatome is not reshaped by a switch from cap-dependent to cap-independent translation, but is instead associated with a preference for GC-rich 3’UTRs and with sequence-specific recruitment of repressive RBPs and miRNAs.

### Senescence collapses a proliferation-specific translational bias favoring transcripts rich in A/U-ending codons

Since codon-level features can shape translation rates, we asked whether specific codons were systematically associated with ΔTE. Fitting a binomial GLM across all expressed coding genes (n = 13,583), we obtained a coefficient for each of the 61 sense codons describing its association with ΔTE (Fig. 5A). The coefficients partitioned almost perfectly by the identity of the codon’s third base: codons ending in G or C were positively associated with ΔTE, while codons ending in A or U were negatively associated. The effect intensified from SEN to LSEN, with LSEN coefficients reaching roughly twice the magnitude of SEN coefficients while largely preserving the same rank order. This effect was not merely due to codon GC content, because the AU/GC axis was specific to the third codon position; positions 1 and 2 showed no comparable bias (Fig. 5A,B). At the transcript level, this manifested as a strong positive relationship between ΔTE and overall third-position GC content (Fig. 5C; SEN R² = 0.19, LSEN R² = 0.30). To control for amino-acid identity, we re-fit the GLM with codon counts normalized to their cognate amino-acid count. This analysis preserved the AU3/GC3 separation (Fig. 5D), confirming that the bias operates at the level of synonymous codon choice rather than amino-acid composition.

**Fig. 5.**
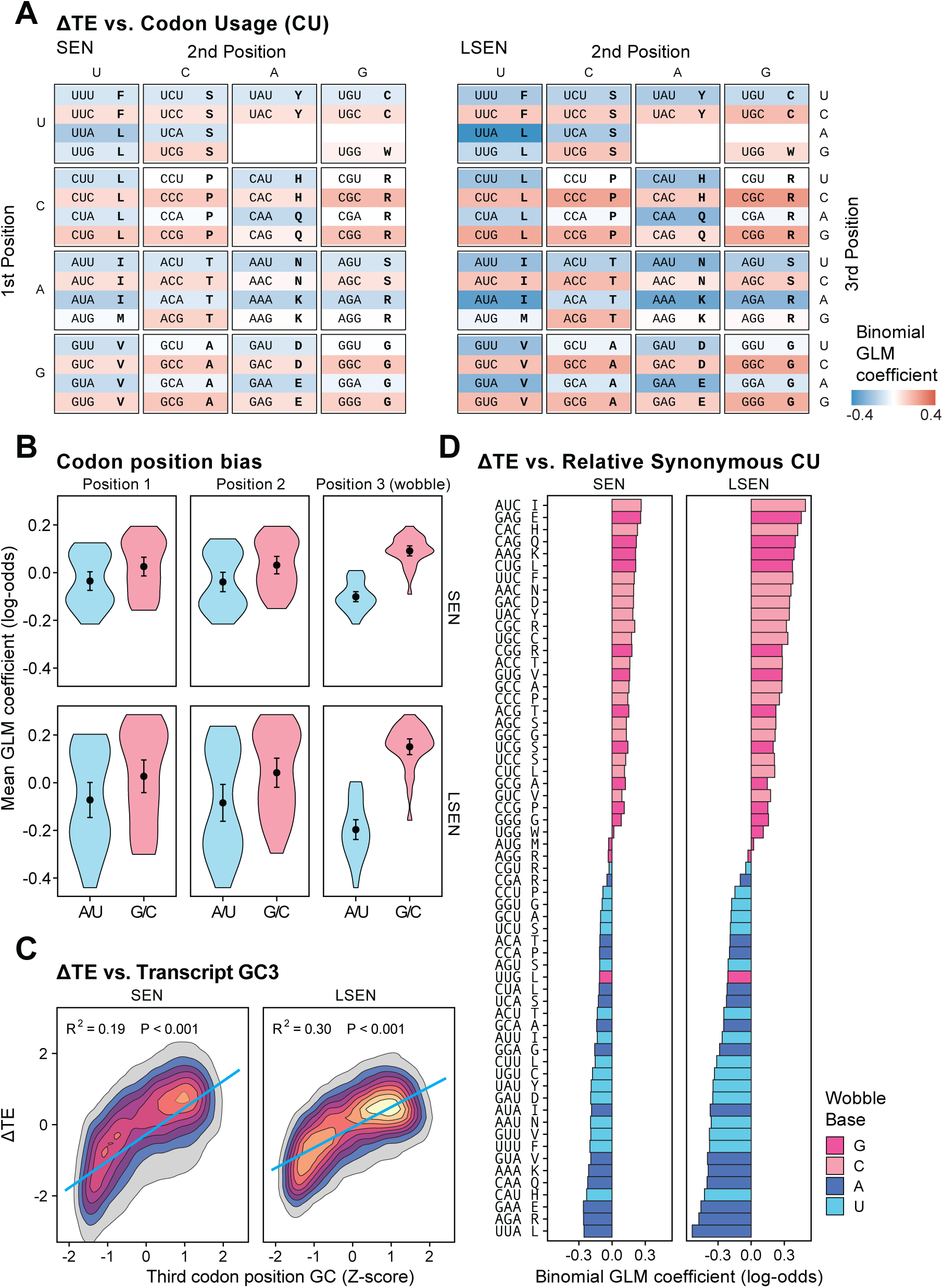
A wobble-position codon bias mediates ΔTE. **A** Per-codon binomial GLMs fit across all expressed coding genes (n = 13,583; average ≥100 counts across conditions). For each of the 61 sense codons, codon proportion was modeled against gene-level ΔTE: (codon count, total codon count − codon count) ∼ ΔTE. Separate models were fit for SEN vs. PRO (left) and LSEN vs. PRO (right). Log-odds coefficients are arranged in the standard codon table and colored by GLM coefficient. **B** Mean per-codon coefficients from (A), stratified by A/U-ending vs. G/C-ending codons at each of the three codon positions, for SEN (top) and LSEN (bottom). Points and error bars: mean ± 95% CI. **C** Per-transcript scatter of ΔTE against transcript wobble-position GC content (GC3 Z-scored across expressed coding transcripts) for SEN (left) and LSEN (right). Density contours overlay individual transcripts; blue line is the linear fit. R² and two-sided p-values for the regression slope are reported. **D** Per-codon binomial GLMs with amino-acid correction—as in (A), but codon occurrence was modeled against cognate amino-acid count: (codon count, amino-acid count − codon count) ∼ ΔTE. Codons are ranked by SEN coefficient and labeled with codon and one-letter amino-acid code; bar fill denotes wobble base.

To understand the origin of this codon bias, we compared translational efficiency (Log2 fold-change of AHA-captured RNA vs. total RNA) to each transcript’s third codon position composition across all conditions (Fig. S8A). In proliferating cells, transcripts enriched in A/U-ending codons were preferentially captured in the translating pool, while those enriched in G/C-ending codons were depleted. This proliferation-specific bias collapsed in both SEN and LSEN, with the regression slope flattening to near zero. The senescence ΔTE signal therefore reflects the loss of an active translation program favoring AU3-rich transcripts in proliferating cells, rather than the emergence of a new program favoring GC3-rich transcripts in senescent cells.

We next asked whether entire gene programs are susceptible to this bias, i.e., whether functional groupings of genes carry sufficiently skewed codon composition to be coordinately affected. We performed gene set enrichment analysis on transcripts ranked by GC3 Z-score (Fig. S8C–E). Among senescence-related gene sets, only the Reactome Cell Cycle set showed a strong association and was sharply AU3-biased (NES = −2.16, padj = 8.6e-16; Fig. S8C,D). Analysis of Hallmark gene sets further identified proliferation-associated sets (Hallmark Myc Targets V1, Hallmark E2F Targets, Hallmark G2M Checkpoint, Hallmark mTORC1 Signaling, Hallmark Mitotic Spindle) as strongly AU3-biased, consistent with previous reports (Gingold et al., 2014). This analysis also revealed that several senescence- and stress-associated programs (Hallmark TNFα Signaling via NF-κB, Hallmark p53 Pathway, Hallmark Hypoxia) were GC3-biased (Fig. S8E). The existence of these biases suggests that the cell can exploit them for regulatory purposes. Notably, while proliferation-associated genes are most strongly enriched for AU3 content, the overall association between ΔTE and third codon position composition does not merely reflect changes in translational efficiency for these genes. We grouped genes into proliferation-related and non-proliferation-related by whether they were nested in at least one of the following GO processes: cell cycle, proliferation, cell division, DNA replication. This partitioning yielded 4,449 “proliferation” genes and 15,018 “non-proliferation” genes, and both sets showed the same negative relationship between GC3 and ΔTE (Fig. S8F). Thus, although cell cycle and other proliferation-related gene sets are disproportionately AU3-rich at the sequence level — and therefore particularly liable to be affected by a change in third codon-position-mediated translational regulation — the overall association of ΔTE to AU/GC3 is not reducible to a translational bias against proliferation-related gene sets.

The mechanisms underlying the observed codon bias are likely multifactorial. Changes in tRNA isoacceptor levels or modification state are two attractive possibilities. The latter can be indirectly assessed from our data using tRNA modification pathway gene expression as a proxy for activity. We examined three wobble-modification systems acting on distinct tRNA subsets: the I34 adenosine deaminases, the Q34 transglycosylases, and the Elongator-mcm⁵U pathway. In each case, a core catalytic or cofactor component was translationally downregulated in senescence (Fig. S8G): ADAT2/3 (I34), QTRT1/2 (Q34), and KTI12, an Elongator cofactor. Translating these changes into precise codon-level predictions is not straightforward, since each modification can affect both A/U- and C/G-ending codons, and the magnitude of the effect at any given codon depends on the availability of competing isoacceptors that can share the decoding load. Nevertheless, these substantial alterations in tRNA anticodon modification pathway expression, which determine codon decoding efficiency, are strong candidate regulators of codon-mediated changes in translational efficiency.

Together, these analyses identify synonymous codon bias as a major axis of translational remodeling in senescence. The proliferating-cell bias toward AU3-rich and against GC3-rich transcripts collapses in senescence. Because cell-cycle and other proliferation gene sets are distinctively AU3-rich, this shift may help mediate and enforce the cessation of proliferation in senescence.

## Discussion

We profiled the translatome of proliferating, senescent, and late-senescent fibroblasts using AHARIBO and focus here on three principal findings. First, we identified a small set of intact L1HS elements that sustain translation across the senescence time course, including one locus whose derepression is supported by chromatin and methylation findings from parallel work in our group. Second, we found that an unexpectedly stringent post-transcriptional depletion of the inflammatory SASP in senescence is relieved as cells progress to late senescence. Third, synonymous codon choice strongly influences changes in translational efficiency in senescence, where we observed the collapse of an AU3-biased translatome. We discuss each finding in turn.

Locus-level identification of actively translated L1 elements has been a persistent obstacle to precisely targeting and studying the L1s that trigger an IFN-I response in senescence. When only mRNA is profiled, it is unclear which, if any, of the expressed L1 elements are being translated: these elements may be sequestered in translationally inactive compartments or otherwise ignored. Ribo-seq approaches digest transcripts into short fragments that are difficult to map uniquely to L1 loci. Proteomics-based approaches are hampered by the near-identity of the vast majority of ionizable L1-derived peptides, as well as by the very low concentrations of ORF2. As a result, the loci producing the inflammatory ORF2 have remained unidentified.

AHARIBO, paired with locus-resolved mapping against a sample-specific reference genome that incorporates non-reference L1 insertions, enabled us to examine the translation of L1s, and RTEs more broadly, in senescence at the level of individual loci. Of all young RTE classes, young L1 elements show the greatest degree of active translation in both SEN and LSEN. This is unsurprising, given that they are the only RTEs that retain a high number of intact open reading frames (Alu and SVA are non-coding, and most ERVs are highly diverged and have largely lost their translational capacities).

Among full-length L1HS loci, those retaining intact ORF1 and ORF2 coding capacity are of particular biological interest, since they have the potential to generate ORF2, and hence trigger cytosolic nucleic acid sensing, cGAS-STING activation, and an IFN-I response. Of these, only two reached significance for upregulation in the AHA translatome at both SEN and LSEN: the intergenic, reference element L1HS_14q23.2_3, and the non-reference element L1HS_3q12.1_1 (Fig. 4D,S4C). The recovery of a non-reference insertion among this short list illustrates the value of building sample-specific reference genomes for studies of young, polymorphic L1 elements; an analysis restricted to the standard human reference genome would have missed it entirely. More broadly, our locus-resolved view paints a dynamic picture of L1 regulation in which only a small subset of intact loci sustains translation across the senescence time course, while others are transiently activated (such as L1HS_1q25.3, which switches on in SEN and off again in LSEN, Fig. 4D). This suggests that cellular defenses against retrotransposition are not disabled in senescence but rather weakened, possibly in a locus-selective manner.

L1HS_14q23.2_3 stands out as a particularly important element. A companion Hi-C study of replicatively senescent LF1 fibroblasts (Dalgarno et al., 2025) identified this same locus as one of just two intact L1HS elements significantly upregulated. Furthermore, it was distinguished from other intact L1HS loci by a striking convergence of activating chromatin features: a shift toward the active A compartment (one of the strongest activating subcompartment transitions), the formation of a senescence-specific chromatin loop placing it in contact with two upregulated genes, and profound demethylation of its promoter. We also observed upregulation of this locus in an independent dataset profiling DNA damage-induced senescence in human astrocytes (Woodham et al., 2026). Our data here show that it is not only transcribed but actively engaged by elongating ribosomes across two senescence time points.

The recurrence of L1HS_14q23.2_3 across three datasets spanning two cell types, two senescence-inducing stimuli, and complementary readouts (chromatin architecture, DNA methylation, transcription, translation) supports a model in which the population of L1HS elements driving inflammatory phenotypes in senescence may be small and architecturally selected, with this locus a leading candidate for functional follow-up.

Beyond L1, AHARIBO revealed translational regulation as a substantive feature of the senescent state. Global transcript levels explained 34% of expression variance in the translatome in SEN, and 70% in LSEN, indicating that translational regulation is a particularly prominent layer of control during the establishment of senescence and becomes less dominant as cells settle into the late state (Fig. 1C,E). Senescent cells have been reported to exhibit a global decrease in translating ribosomes (Payea et al., 2024), which our data corroborate (Fig. S1A); we show, however, that they do not merely undergo a uniform translational decline but actively regulate key senescence programs at the translational level: cell-cycle genes are translationally suppressed, while extracellular-matrix remodeling genes are translationally enhanced (Fig. 2B,C).

In our data, the SASP provided a striking instance of post-transcriptional regulation: under strong translational control, it exhibited a dynamic phenotype across the senescence time course, translationally depleted in SEN and derepressed in LSEN (Fig. 3). Motif analysis nominated the ZFP36 family alongside a broader cohort of AU-rich element (ARE)-binding proteins as key candidate regulators of this effect (Fig. 3C,D), consistent with the established role of ZFP36L1 in SASP regulation in oncogene-induced senescence (Herranz et al., 2015). However, the strong inter-factor correlation among ARE-binding motifs (Fig. S3) means this analysis cannot resolve which factor or factors mediate the observed regulation, and orthogonal approaches such as CLIP or RBP-specific knockdown ribo-seq will be required to validate and deconvolve their relative contributions.

To investigate the mechanisms underlying translational-efficiency changes in senescence, we correlated coding gene ΔTE with various transcript sequence features and motifs (Fig. 5, S6). The strongest sequence-level determinant of ΔTE was codon third-position GC content: in proliferating cells, AU3-rich transcripts are translationally favored and GC3-rich transcripts disfavored, and this preference is lost in senescence (Fig. 5A,B; S8A). Proliferation-associated gene sets are distinctively AU3-rich, as first noted by Gingold et al. (Gingold et al., 2014), and our GSEA recovers cell-cycle and proliferation programs (Reactome Cell Cycle, Hallmark Myc Targets V1, Hallmark E2F Targets, Hallmark G2M Checkpoint, Hallmark mTORC1 Signaling, Hallmark Mitotic Spindle) as the strongest AU3-biased sets in the genome (Fig. S8C–E). However, the ΔTE–GC3 relationship is not reducible to these gene programs: partitioning genes by membership in proliferation-related GO terms revealed the same negative ΔTE–GC3 relationship in both groups (Fig. S8F). The codon bias therefore operates as a genome-wide translational filter, with proliferation-related genes disproportionately affected owing to their AU3-rich architecture rather than serving as the sole substrate of the effect. Our data extend Gingold’s dual-program framework with direct translatome evidence that the AU3-favoring translation program is actively withdrawn as cells exit proliferation.

A mechanistic candidate for the loss of AU3-favored translation in senescence is modulation of tRNA wobble-modification machinery. ADAT2/3, QTRT1/2, and KTI12 (an Elongator cofactor) were all translationally downregulated in both SEN and LSEN. This parallels prior work in pluripotent stem cells, where ADAT2 declines as ESCs exit self-renewal with concomitant loss of I34 on ADAT-substrate tRNAs (Bornelöv, Selmi, Flad, Dietmann, & Frye, 2019); conversely, multiple wobble-modification enzymes are recurrently upregulated or amplified in human malignancies (Ramirez-Moya et al., 2025; Rapino, Delaunay, Zhou, Chariot, & Close, 2017). Whether these alterations in tRNA modification pathways cause the senescence codon bias is difficult to establish from expression data alone: each modification can influence both A/U- and C/G-ending codons depending on context. Direct measurement of tRNA modification states and codon-resolved ribosome dwell times will be required to formally test this.

Together, these findings establish translational control as a substantive regulatory layer in cellular senescence, governing both the SASP and locus-selective protein expression of intact L1 retrotransposons, and disfavoring proliferation-associated genes through biased translation at the level of synonymous codon choice, possibly mediated by the loss of tRNA wobble modifications. Notably, among intact L1HS loci, L1HS_14q23.2_3 stands out: it is one of only two intact elements with sustained translation across both senescence states in this study, and one of only two significantly upregulated in parallel work from our group, where it was further distinguished by uniquely strong activating chromatin and methylation changes.

## Methods

### Cell Culture and Senescence Establishment

Human fetal lung fibroblast (LF1) cells were recovered from frozen stocks and expanded for experimentation (Brown, Wei, & Sedivy, 1997). Cells were maintained at 37 °C in a humidified incubator under physiological oxygen (2.5% O₂, 5% CO₂). Cultures were grown in Ham’s F-10 nutrient mixture (Gibco) supplemented with 15% fetal bovine serum (FBS; Corning), 2 mM L-glutamine, and penicillin–streptomycin. Mycoplasma contamination was routinely monitored with the MycoAlert Mycoplasma Detection Kit (Lonza).

Proliferating (PRO) cells were obtained from early-passage cultures. Replicative senescence was induced by serial passaging until cell division ceased. At each passage, cultures were grown to ∼80% confluency, trypsinized, and diluted 1:4. In early-passage cultures, the interval between passages was ∼4–5 days, but this progressively extended to 2–3 weeks in the final few passages before senescence. Entry into senescence (t=0) was defined as the date of the last 1:2 passage after which cultures never reached 80% confluency again (see De Cecco et al. 2019 for a more complete description). Senescent cells (SEN) were harvested 4-8 weeks post t=0, and late-senescent cells (LSEN) were harvested 16 weeks post t=0.

For the PRO state, a single 80% confluent 10-cm dish yielded sufficient material for RNA isolation. By contrast, SEN and LSEN cells required 3-5 10-cm dishes to obtain comparable yields. This difference reflects distinct cell morphologies: proliferating cells are small with the typical elongated fibroblastic shape, whereas senescent cells are much larger and flatter, lowering cell density per dish.

### Total RNA Isolation

PRO, SEN, and LSEN LF1 cells were cultured in 10-cm dishes. At ∼80% confluency, culture medium was aspirated, and cells were lysed by adding 1 mL TRIzol reagent directly to the dish (on ice). Plates were swirled to ensure complete coverage and scraped with a sterile cell scraper. Lysates were collected into 15-mL conical tubes and incubated at room temperature for 5 min. Chloroform (200 µL) was added, samples were vortexed briefly, incubated for 2 min, and centrifuged at >12,000 × g for 15 min at 4 °C. The aqueous phase was transferred to a fresh tube, mixed with an equal volume of freshly prepared 70% ethanol, and applied to an RNeasy Mini spin column (Qiagen). RNA was purified according to the manufacturer’s instructions and eluted in RNase-free water. This method recovers all RNA species >200 nucleotides.

### AHARIBO Isolation of Actively Translated RNA

The AHARIBO assay (IMMAGINA BioTechnology, cat. no. AHA003) was used to isolate actively translating ribosome complexes by metabolic labeling with L-azidohomoalanine (AHA), a non-toxic methionine analog incorporated into nascent peptides during protein synthesis. PRO, SEN, and LSEN LF1 cells were grown to ∼80% confluency in 10-cm dishes. Cells were washed once with PBS and incubated with methionine-free medium (Thermo Scientific, cat. no. 30030) for 40 min at 37 °C. AHA (1 mM final concentration) was added, and cells were incubated for 5 min at 37 °C, followed by treatment with sBlock (IMMAGINA BioTechnology, cat. no. RM8; a proprietary anisomycin-containing reagent that blocks nascent peptides on ribosomes) for 5 min at 37 °C. Plates were placed on ice, washed with ice-cold PBS, and lysed in AHARIBO lysis buffer using a cell scraper. Lysates were clarified by centrifugation at 20,000 × g for 5 min at 4 °C, transferred to new tubes, and kept on ice for 10 min. Nucleic acid concentration was measured at 260 nm with a NanoDrop spectrophotometer, using lysis buffer as the blank.

Clarified lysates corresponding to 0.2 absorbance units (260 nm) were supplemented with W-buffer containing RiboLock. Functionalized sBeads were added, and samples were incubated for 60 min at 4 °C on a rotating wheel (9 rpm). Beads were collected on a magnetic rack, washed with WSS solution (10 min, 4 °C, 9 rpm), and resuspended in W-buffer. The bead suspension, containing ribosome–RNA complexes, was transferred to fresh nuclease-free tubes. RNA was extracted from the ribosome complexes using the AHARIBO protocol according to the manufacturer’s instructions and further purified by the acid phenol–chloroform method. RNA quality and quantity were assessed using an Agilent TapeStation.

### RNA-seq and Library Preparation

Total RNA was extracted from PRO, SEN, and LSEN cells as described above. Polyadenylated RNA was enriched using the NEBNext Poly(A) mRNA Magnetic Isolation Module (NEB #E7490), and strand-specific cDNA libraries were prepared with the NEBNext Ultra II Directional RNA Library Prep Kit for Illumina (NEB #E7765S/L), following the manufacturer’s protocol. Library quality and concentration were assessed using an Agilent TapeStation. Validated libraries were sequenced by Azenta using paired-end reads on an Illumina NovaSeq 6000 platform. Four biological replicates were processed per condition.

### RNA-seq core analysis

Samples were comparable in read quality and mapping scores, and were deeply sequenced, with library size averaging 128M reads (Table S1). We analyzed these data with our TE-Seq pipeline and custom scripts. Full TE-Seq pipeline methods can be found at https://doi.org/10.1186/s13100-025-00381-w. Briefly, sequencing reads were trimmed with Fastp and aligned to a sample-specific TE-patched genome (see Methods: Non-reference Germline Insertion Calling and TE-Patching Sample Genomes) with STAR. Gene expression was quantified using featureCounts and a RefSeq transcript annotation (GCF_009914755.1). Repetitive element expression was quantified using the telescope tool in both “unique” and “multi” modes, which considered either uniquely mapped reads or both uniquely and multi-mapped reads.

Genes were assessed for differential expression using DESeq2. When comparing expression within a library type (e.g. AHA SEN vs. AHA PRO), we used median of ratios normalization size factors. When comparing expression across library type (e.g. AHA SEN vs. TOT SEN), we used size factors derived from the total number of mapped and paired reads, since the median of ratios assumption that most genes have the same expression between conditions is likely violated when comparing the translatome to the transcriptome. As our measure of translational efficiency, TE, we used the Log2 fold-change between AHA and TOT counts for a given biological condition. As our measure of ΔTE, we used the difference between the AHA Log2 fold-change and TOT Log2 fold-change between biological conditions.

We used GSEA to query MSigDB gene sets for enrichment, ranking genes by Log2 fold-change or by the change in translational efficiency, ΔTE. Genome tracks were plotted using Plotgardener. Differential expression analysis of aggregate RTE subfamily counts was performed using a negative binomial GLM. For this analysis, raw counts were summed across group members, and sample size factors were incorporated as an offset in the model.

### Transcript RBP motif analysis

To identify putative RNA-binding protein translational regulators, we used the Gimme Motifs suite (van Heeringen & Veenstra, 2011). UTR sequences of SenMayo SASP genes were scanned for RNA-binding protein motifs from the CIS-BP database using Gimme Motifs “gimme scan” in sense-strand mode (--norc), with per-motif score thresholds calibrated at a 5% false-positive rate against GC- and length-matched random genomic regions, reporting one hit per transcript per motif (--table --nreport 1 -f 0.05). For each RBP, ΔTE was regressed on binary motif presence using a univariate linear model. Additionally, an elastic net model was fit on all RBPs simultaneously.

We used Gimme Motifs “gimme maelstrom” to search for motifs enriched in translationally enhanced or depleted coding genes. We partitioned the UTRs of all protein-coding genes into discrete classes — enriched (ΔTE > 2), depleted (ΔTE < −2), and non-changing (0.2 > ΔTE > −0.2) — and searched for motifs comparing enriched and depleted to non-changing, or enriched to depleted classes. RBP motifs from the CIS-BP database were used. Analyses were run both with and without maelstrom’s GC-content correction (--nogc).

### Regulatory Module classification

We classified genes into distinct regulatory modules based on the directionality of their transcriptional changes (ΔTx) and changes in their translational efficiency (ΔTE). We required transcriptional changes to be called significant by DESeq2 and to have a Log2FC ≥ 0.585, and considered genes with translatomic changes that differed by at least a factor of 2 from transcriptomic changes (ΔTE ≥ 1; ΔTE is on the Log2 scale) to be post-transcriptionally regulated in senescence.

### Codon usage and wobble-position analyses

Per-gene codon frequencies for human RefSeq Select protein-coding transcripts were obtained from the CoCoPUT database (Athey et al., 2017). For each gene, we computed counts of the 61 sense codons, the cognate amino-acid counts, and wobble-position GC content (GC3, the fraction of codons ending in G or C), Z-scored across expressed coding genes (defined as genes with an average of ≥100 counts across conditions; n = 13,583).

Per-codon binomial GLMs were fit with glmmTMB (Brooks et al., 2017), modeling per-gene codon proportion against ΔTE — the difference between the AHA and TOT Log2 fold-changes for a given senescence vs. PRO comparison. In the unadjusted model, the denominator was total codon count: glmmTMB(cbind(codon_count, total_codon_count − codon_count) ∼ ΔTE, family = binomial). In the amino-acid-adjusted model, the denominator was the cognate amino-acid count. The unadjusted model is required for positions 1 and 2 because amino-acid normalization would confound any GC1/GC2 signal with amino-acid usage; the adjusted model isolates the synonymous-codon contribution of position 3.

For per-transcript regressions, ΔTE (or TE, the AHA vs. TOT Log2 fold-change within a single condition) was regressed on the GC3 Z-score by ordinary least squares. For proliferation-stratified analyses, genes were partitioned by membership in the GO Biological Process terms cell cycle, cell proliferation, cell division, or DNA replication (n = 4,449 proliferation-associated, n = 15,018 other). GC3-ranked GSEA was performed with fgsea against curated senescence-related sets and the MSigDB Hallmark collection.

## Declarations

## Ethics approval and consent to participate

Not applicable

## Consent for publication

Not applicable

## Availability of data and materials

A Zenodo archive of all code used can be found at: 10.5281/zenodo.20434425. The most up-to-date version of the TE-Seq pipeline (which was used for sample-specific, non-reference TE-patched genome generation and RNA-seq analysis in (Dalgarno et al., 2025)) can be found at https://github.com/maxfieldk/TE-Seq, where instructions for pipeline installation, configuration, and deployment are documented. All sequencing data used have been deposited in the Sequence Read Archive (SRA); RNA-seq data: BioProject accession number PRJNA1178097, Nanopore DNA-sequencing data: BioProject accession number PRJNA1375768. Any other data or information relevant to this study are available from the corresponding author upon reasonable request.

## Authors’ contributions

Conceptualization: JMS

Data curation: RLK

Formal analysis: MMK, RLK

Funding acquisition: JMS

Investigation: MMK, RLK, JMS

Methodology: JMS, MMK, RLK

Project administration: JMS

Resources: JMS

Software: MMK

Supervision: JMS

Writing – original draft: All authors

Writing—review & editing: All authors

## Acknowledgments

We would like to thank Jess Anderson for lab management and technical assistance, and members of the Sedivy lab and the Center on the Biology of Aging for feedback and support. We are grateful to the Brown Center for Computation and Visualization (CCV) team for their management of the OSCAR high performance computing cluster which was used throughout this work.

---

**Fig. S1.**
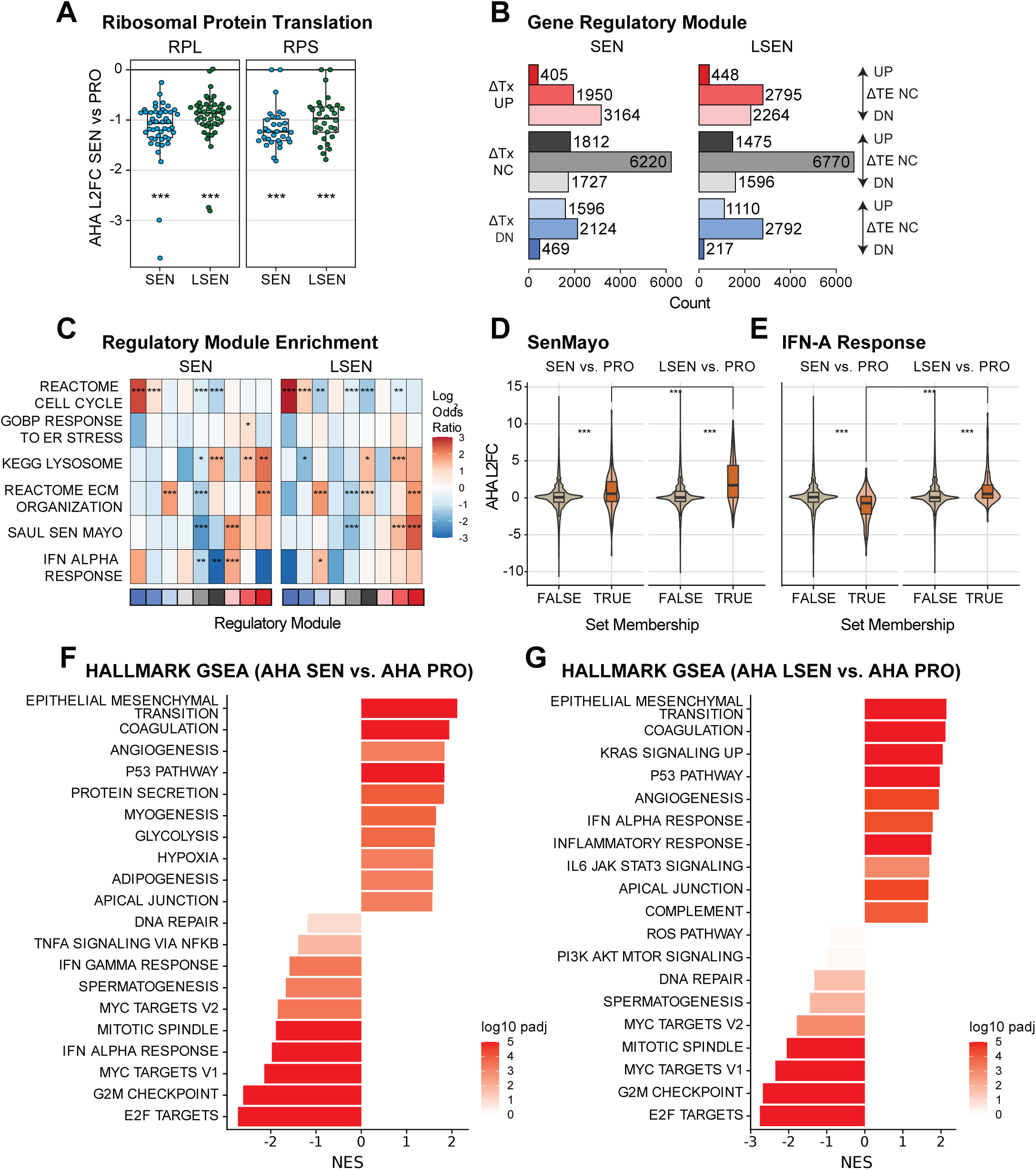
The translatomic landscape of senescence reveals extensive and stage-dependent translational regulation. **A** Translatomic expression Log2 fold-change for ribosomal protein genes (left, large subunit ribosomal proteins, RPL; right, small subunit ribosomal proteins, RPS). Statistical significance of RPL and RPS gene set translational depletion was determined by GSEA against all coding genes. **B** Classification of genes into regulatory modules based on transcriptional (ΔTx) and translational (ΔTE) changes for SEN (left) and LSEN (right). **C** Heatmap showing the enrichment (Log2 Odds Ratio) of senescence-related gene sets for regulatory module genes. Statistical significance is indicated by asterisks (* p ≤ 0.05, ** p ≤ 0.01, *** p ≤ 0.001). **D,E** Violin and box plots of coding gene AHA Log2FC SEN vs. PRO (left) and AHA Log2FC LSEN vs. PRO (right) stratified by membership in the SenMayo gene set (D) or the Hallmark Interferon Alpha Response gene set (E). For all comparisons, statistical significance was assessed with a two-sided Wilcoxon rank-sum test; *** p ≤ 0.001. **F,G** Gene Set Enrichment Analysis (GSEA) of Hallmark gene sets. Genes were ranked by their DESeq2 Wald statistic for the AHA SEN vs. PRO (F) and AHA LSEN vs. PRO (G) comparisons. Bar plots show Normalized Enrichment Scores (NES) and are colored by FDR-adjusted p-value.

**Fig. S2.**
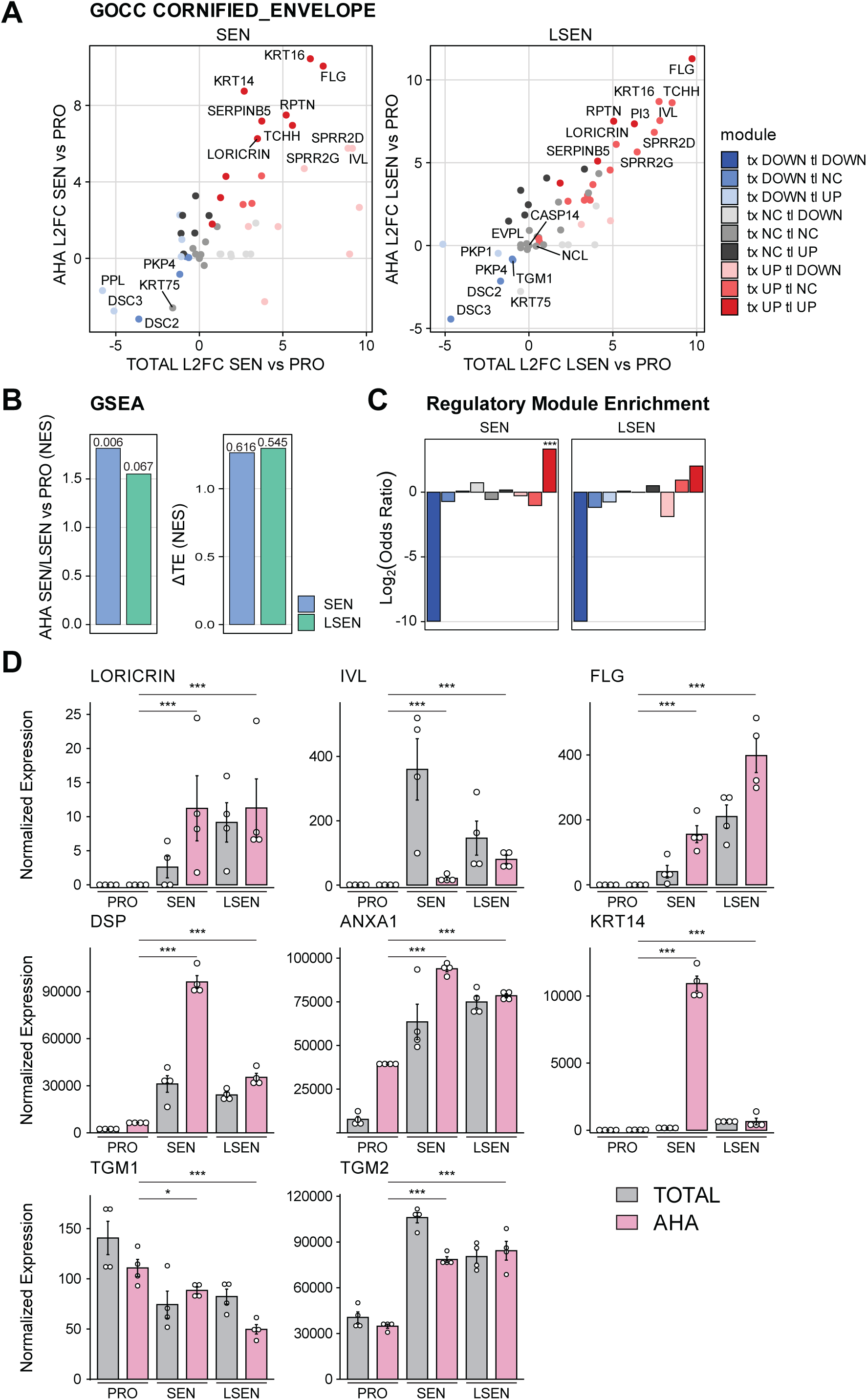
Cornified Envelope (CE) **A** Scatter plots comparing the expression changes of GO CC Cornified Envelope (CE) genes in the translatome vs. the transcriptome for SEN vs. PRO (left) and LSEN vs. PRO (right). **B** GO CC CE gene set GSEA results using genes ranked by (left) AHA Log2 fold-change (SEN/LSEN vs. PRO) or (right) ΔTE. **C** Bar plots displaying the regulatory module enrichment (Log2 Odds Ratio) of the GO CC CE gene set. **D** Gene expression plots for selected CE genes. Error bars represent mean ± SEM. Statistical significance was assessed using the DESeq2 FDR-adjusted p-value. Significance is indicated by asterisks (p ≤ 0.05 *, ≤ 0.01 **, and ≤0.001 ***).

**Fig. S3.**
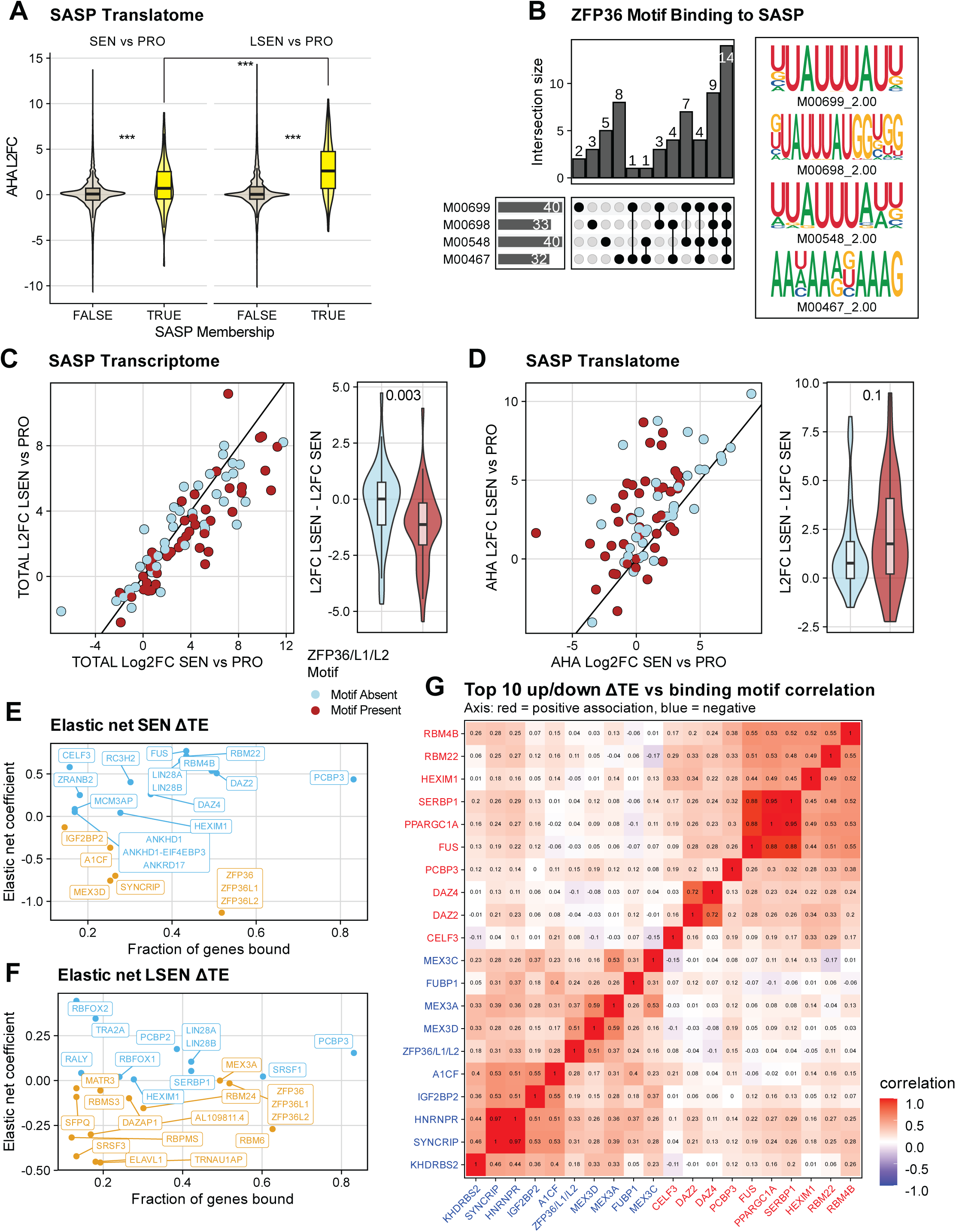
ZFP36-motif-bearing SASP transcripts shift from translational repression in SEN to relief of repression and re-engagement of decay in LSEN. **A** Violin and box plots of coding gene AHA Log2FC SEN vs. PRO (left) and AHA Log2FC LSEN vs. PRO (right) stratified by membership in the SenMayo SASP gene set. SASP transcripts were upregulated in the translatome in both SEN and LSEN, though considerably more so in the latter (SEN median Log2FC = 0.71, LSEN median Log2FC = 2.61). For all comparisons, statistical significance was assessed with a two-sided Wilcoxon rank-sum test; *** p ≤ 0.001. **B** UpSet plot showing the intersection pattern of the four CIS-BP ZFP36-family motifs (M00699, M00698, M00548, M00467) across SenMayo SASP transcripts. Bar heights give the number of transcripts containing each indicated motif combination; row totals (right of motif labels) give the number of transcripts containing each motif individually. Sequence logos for the four motifs are shown at right. **C,D** Scatter plot (left) of SenMayo SASP gene expression in TOTAL RNA-seq, comparing LSEN vs. PRO Log2 fold-change (y-axis) to SEN vs. PRO Log2 fold-change (x-axis). Points are colored by ZFP36/ZFP36L1/ZFP36L2 binding motif presence (present, red; absent, light blue). Violin and box plots (right) of TOTAL Log2FC LSEN vs. PRO minus Log2FC SEN vs. PRO, stratified by ZFP36/ZFP36L1/ZFP36L2 binding motif presence. Statistical significance was assessed with a two-sided Wilcoxon rank-sum test. **D** As in C, for AHA (translatome). **E,F** SenMayo SASP genes were assayed for ZFP36 motif presence using Gimme Motifs. SEN (E) and LSEN (F) ΔTE was regressed against motif presence using an elastic net model. The RBPs with the top regression coefficients are shown, along with the fraction of SASP genes carrying their binding motif. **G** Heatmap of pairwise Pearson correlations between binary motif-presence vectors across SenMayo SASP transcripts, restricted to the ten RBPs with the most positive (red labels) and the ten with the most negative (blue labels) univariate associations with LSEN ΔTE.

**Fig. S4.**
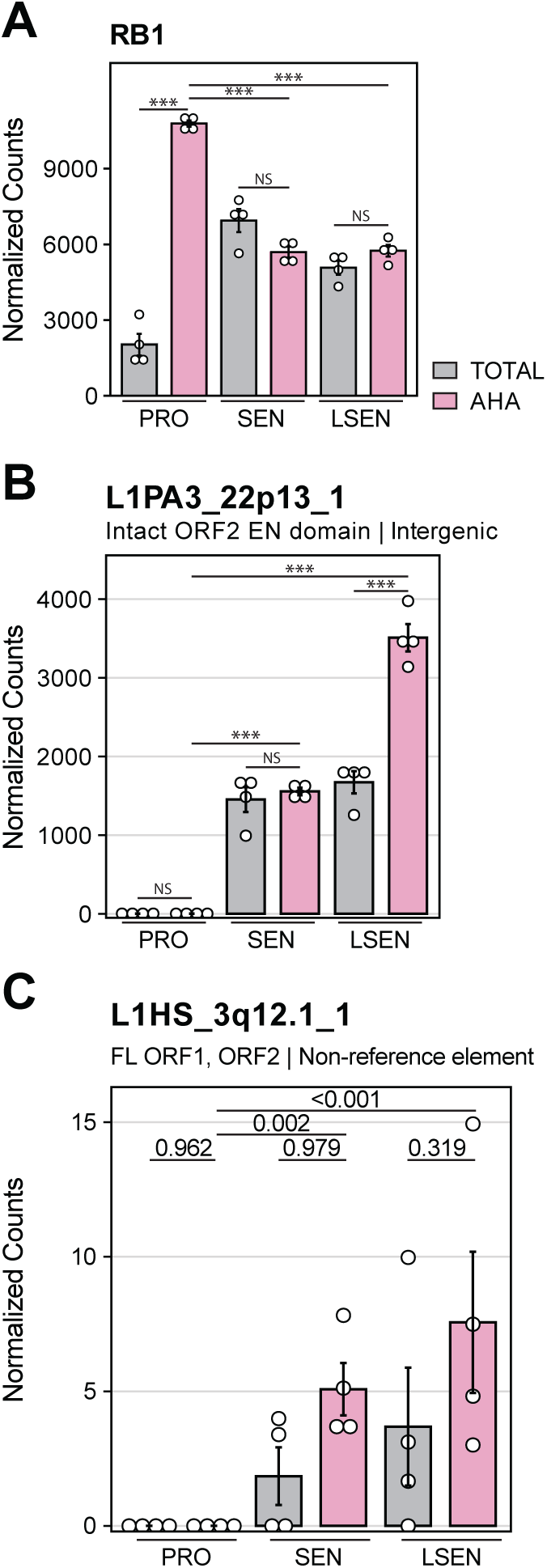
**A** RB1 gene expression. **B,C** Unique count expression profiles for L1PA3_22p13_1 (B) and L1HS_3q12.1_1 (C). Error bars represent mean ± SEM. DESeq2 FDR-adjusted p-values for statistical comparisons between AHA and TOT are shown for each senescence condition, as well as for the AHA SEN/LSEN vs. PRO comparisons.

**Fig. S5.**
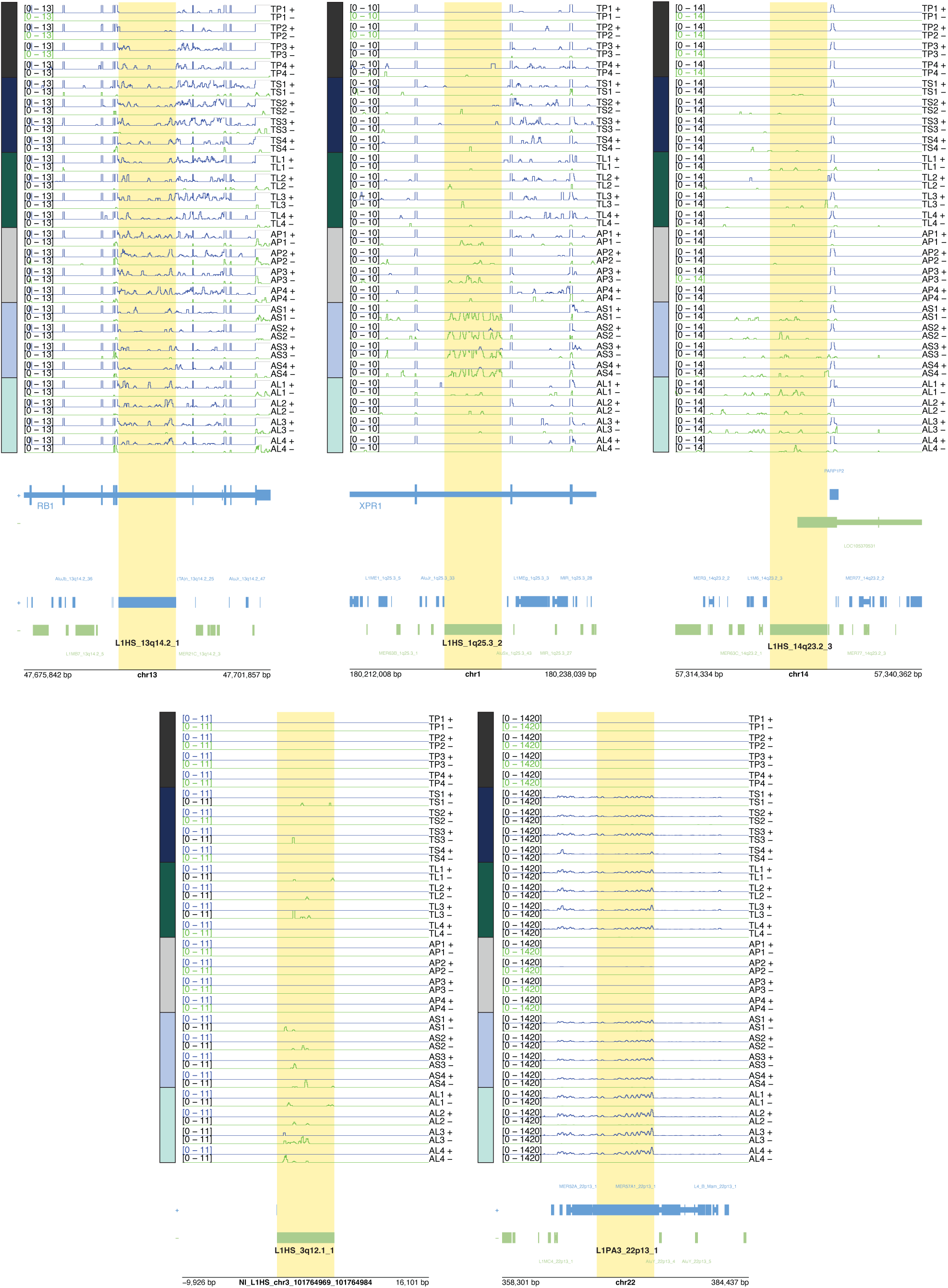
Genome tracks of differentially expressed young L1 elements. Genome tracks of the three differentially expressed, full-length L1HS elements highlighted in Fig. 4C,D, as well as the differentially expressed non-reference element L1HS_3q12.1_1. Tracks for the markedly upregulated L1PA3_22p13_1 are also shown. Unique alignments (non-unique alignments are not shown) were converted to BigWig format and visualized in R using Plotgardener. BigWig tracks of Watson (+; blue) and Crick (-; green) strand transcription are plotted for each sample, followed by RefSeq and RepeatMasker annotations. The region is centered on the L1HS element (boxed in black), with 10 kb of flanking up/downstream sequence. The BigWig signal track’s Y-axis upper bound is set at 110% of the maximum count value observed across all samples over the L1HS locus.

**Fig. S6.**
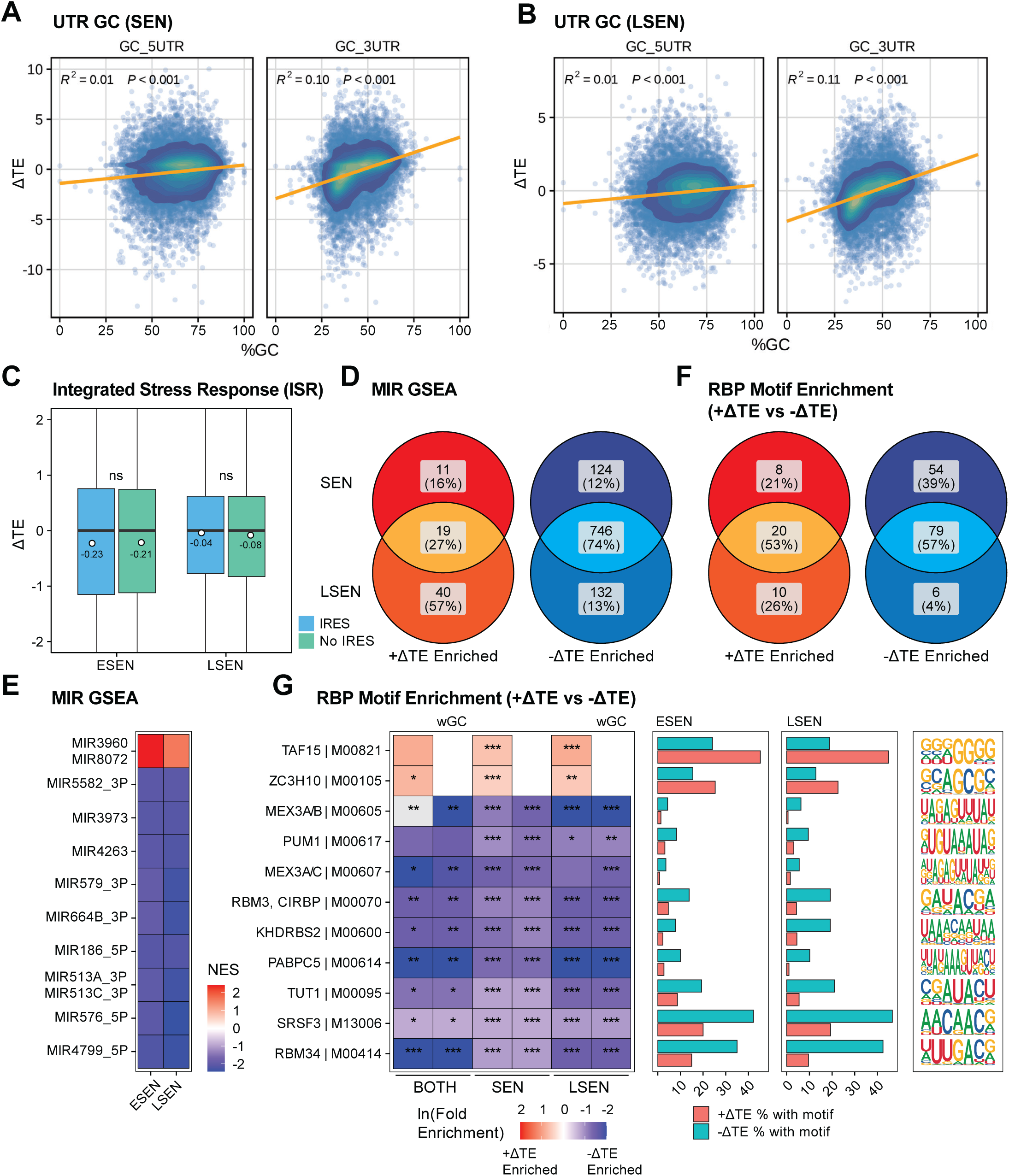
Sequence features shape translational efficiency in senescence. **A,B** Scatter plots illustrating the relationship between ΔTE and GC content (%) of the 5’ Untranslated Region (5’UTR, left) and 3’ Untranslated Region (3’UTR, right). Relationships are shown for (A) SEN and (B) LSEN conditions. A positive correlation (R² ≈ 0.10-0.11, p ≤ 0.001) is observed specifically between ΔTE and 3’UTR GC content, indicating that GC-rich 3’UTRs are preferentially translated in senescence. Color intensity represents data point density. **C** Boxplot of ΔTE for internal ribosome entry site (IRES)-containing and non-IRES-containing genes (as annotated by IRESbase[42]). Statistical significance was assessed with a two-sided Wilcoxon rank-sum test. **D,F** Venn diagrams showing the number of microRNA (miRNA)-regulated gene sets (D) and RNA-binding protein (RBP) motifs (F) associated with translational enhancement or depletion. RBP motif enrichment within transcript UTRs was assessed using Gimme Motifs with and without GC-correction (for GC-corrected Venn diagrams, see Fig. S7A). Genes with an average of at least 100 counts in one condition that exhibited translational enrichment were compared to those exhibiting translational depletion (ΔTE ≥ 2 vs. ΔTE ≤ −2). miRNA gene set enrichment was determined by GSEA. **E** Heatmap of miRNA-regulated gene set GSEA for genes ranked by ΔTE. Normalized Enrichment Scores (NES) are shown. **G** RNA-binding protein (RBP) motif enrichment analysis: (left) heatmap showing the natural log fold motif enrichment for +ΔTE (red) and −ΔTE (blue) genes. Statistical significance was assessed using a hypergeometric distribution on motif occurrence counts, with Benjamini-Hochberg-adjusted p-values reported, p ≤ 0.05 *, ≤ 0.01 **, and ≤ 0.001 ***. (middle) Bar plots show the percentage of target transcripts containing the motif. (right) Logos depict the consensus recognition sequences.

**Fig. S7.**
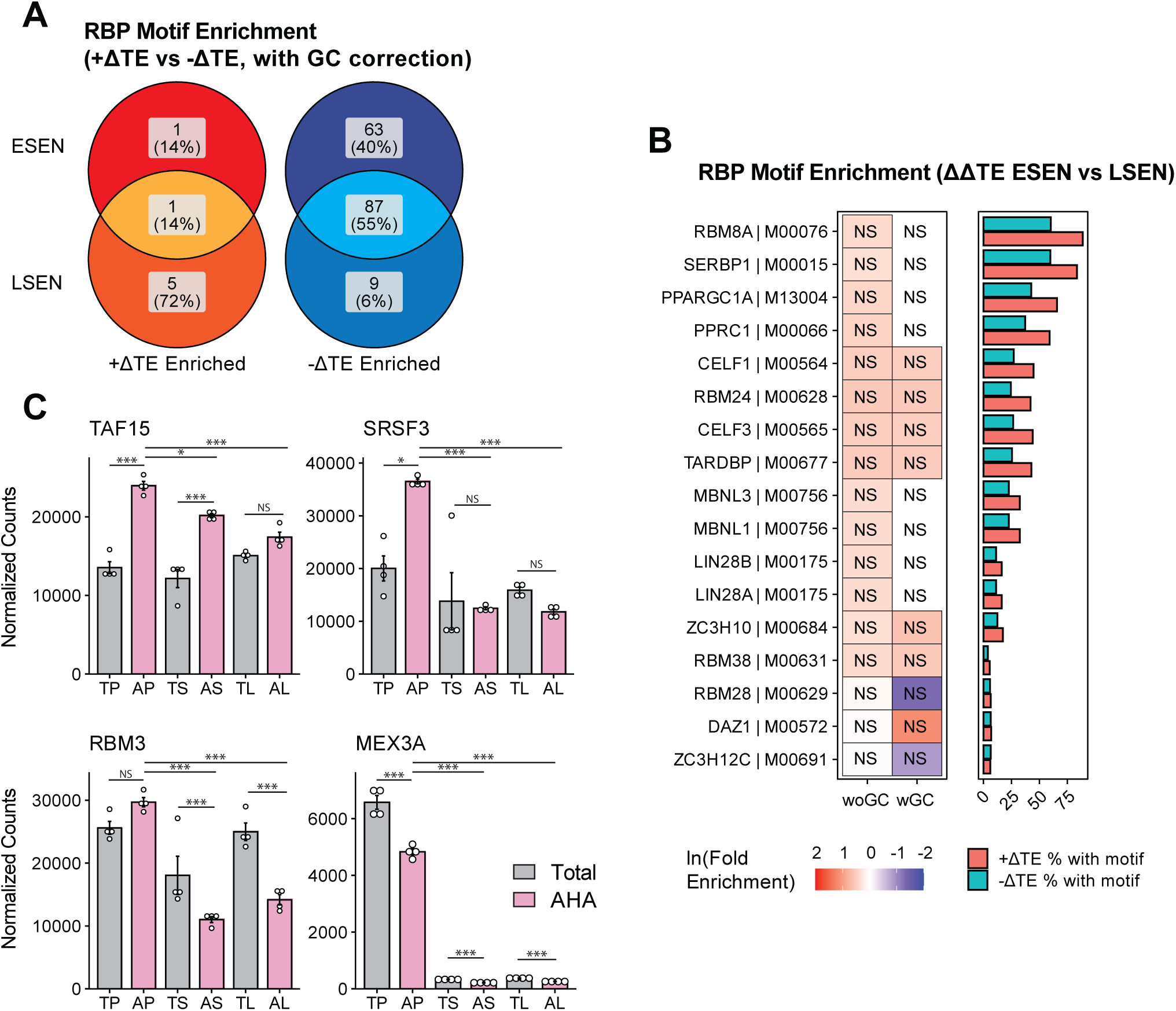
Sequence features shape translational efficiency in senescence, continued. **A** Venn diagram showing the number of RNA-binding protein (RBP) motifs associated with translational enhancement or depletion after GC content correction (for non-GC corrected values, see Fig. S6G). **B** RNA-binding protein (RBP) motif enrichment analysis (using Gimme Motifs) of transcripts exhibiting variable translational regulation between SEN and LSEN (|ΔΔTE | ≥ 2). (left) Heatmap showing the natural log fold motif enrichment for +ΔΔTE (red, higher ΔTE in SEN) and −ΔΔTE (blue, higher ΔTE in LSEN) genes. Statistical significance was assessed using a hypergeometric distribution on motif occurrence counts. (middle) Bar plots show the percentage of target transcripts containing the motif. **C** Gene expression plots for select top enriched RBPs in ΔTE SEN/LSEN vs. PRO (Fig. S6G).

**Fig. S8.**
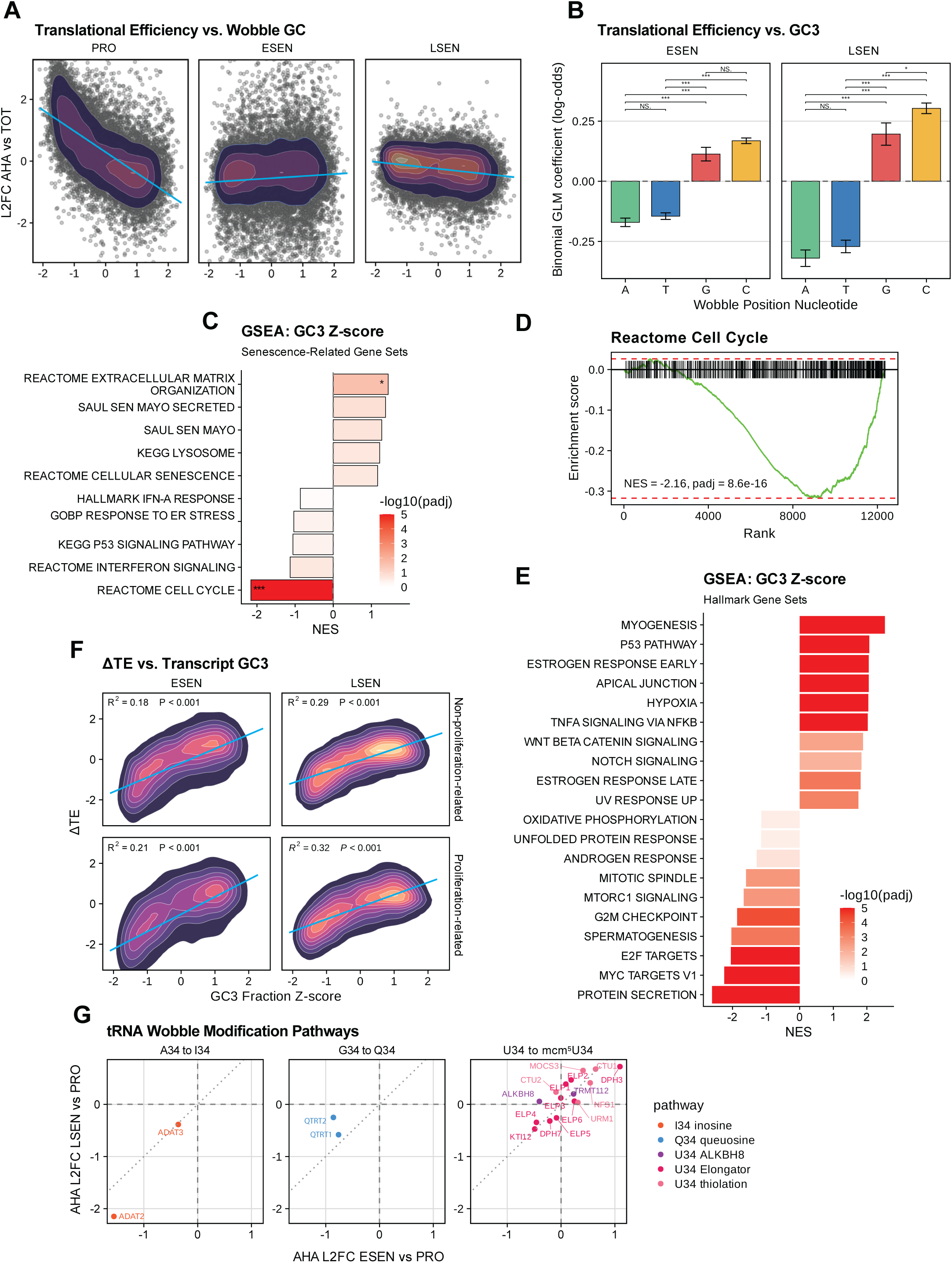
Wobble-position codon bias effect on ΔTE reflects the loss of an AU3-favoring translation program operant in proliferating cells. **A** Per-transcript scatter of translational efficiency (AHA Log2 fold-change vs. TOT Log2 fold-change) against wobble-position GC content (GC3 Z-scored across expressed coding transcripts), for PRO, SEN, and LSEN. Density contours overlay individual transcripts; the blue line is the linear regression. **B** Per-codon binomial GLM coefficients from Fig. 5A, stratified by wobble-position nucleotide (A, U, G, C) for SEN (left) and LSEN (right). Bars: mean ± 95% CI. Significance from pairwise comparisons: NS, not significant; *p ≤ 0.05; ***p ≤ 0.001. **C** GSEA of expressed coding genes ranked by GC3 Z-score, tested against selected senescence-related gene sets. NES, normalized enrichment score; bar fill denotes −log10(padj). *FDR ≤ 0.05, **≤ 0.01, ***≤ 0.001. **D** Enrichment plot for Reactome Cell Cycle from the analysis in (C). **E** As in (C), tested against the MSigDB Hallmark gene sets. **F** Per-transcript scatter of ΔTE against transcript wobble-position GC content (GC3 Z-scored across expressed coding transcripts) for SEN (left) and LSEN (right), stratified by membership in proliferation-associated GO processes – Cell Cycle, Cell Proliferation, Cell Division, DNA Replication. Density contours overlay individual transcripts; the blue line is the linear regression. R² and two-sided p-values for the regression slope are reported. **G** Translatomic expression changes of tRNA wobble-modification enzymes, grouped by pathway. AHA Log2 fold-change LSEN vs. PRO plotted against AHA Log2 fold-change SEN vs. PRO for each modification system; A34→I34 inosine deamination (top), G34→Q34 queuosine incorporation (middle), and U34→mcm⁵U34 or related modifications (decomposed into ALKBH8 methylation, Elongator, and thiolation sub-pathways).

